# The DLX/Notch axis is necessary for spatiotemporal regulation of neural cell fate

**DOI:** 10.1101/2025.09.28.679022

**Authors:** Ryan F. Leung, Michael See, Ankita M. George, Patrick L. Barry, Natalie Charitakis, Mirana Ramialison, Maree C. Faux, David D. Eisenstat

## Abstract

The neuronal-glial cell fate switch during forebrain development is highly regulated. DLX transcription factors are necessary for promoting GABAergic interneuron differentiation and migration but the mechanisms for concomitant repression of glial fate in neural progenitors remain elusive. Here, the DLX2 regulatory network dynamic in the developing ventral telencephalon was characterised using a multi-omic approach at single-cell resolution, including single-cell whole genome spatial transcriptomics. We identified a secondary proliferative zone in the ventral subventricular zone and spatiotemporal-context dependent Notch pathway repression by DLX2 in maintaining progenitor populations and facilitating neural differentiation. We found that DLX2 controls cell fate determination by directly repressing Notch signalling genes as well as glial fate promoting transcription factors, thereby inhibiting early adoption of oligodendroglial differentiation during neurogenesis. Thus, temporal cell fate switch mediated by DLX2 via a multilayer gene regulatory network redefines our current understanding of neuronal-glial cell specification mechanisms in the developing telencephalon.

## Introduction

The developing forebrain includes the telencephalon and diencephalon. The telencephalon is primarily comprised of the neocortex in the pallium, and the ganglionic eminences (GE) in the subpallium. In mice, neurogenesis begins in the GE at E10.5, followed by overlapping waves of oligodendrogenesis and astrocytogenesis (Cadwell et al., 2019). Neural progenitors in the GE significantly contribute to γ-aminobutyric acid (GABA)-ergic interneurons (INs), making up over two-thirds of cortical INs in mice (Anderson et al., 2002).

IN differentiation and migration are regulated by key signalling pathways and transcription factors (TFs), including the *Distalless homeobox (Dlx)* TFs (Anderson, Eisenstat, et al., 1997; Le et al., 2017). DLX TFs can function as activators or repressors, and a recent study showed transcriptomic consequences of DLX binding can be predicted by the epigenomic context of its target loci (Lindtner et al., 2019). DLX TFs promote IN migration via signalling molecules (*Cxcr4*, *Nrp2*, *Pak3*) (Cobos et al., 2007; Le et al., 2007; Wang et al., 2011), evident through the loss of their tangential migration in *Dlx1/Dlx2* double knockout (*Dlx1/Dlx2^-/-^*) mouse GE(Anderson, Eisenstat, et al., 1997). DLX2 also promotes IN differentiation via promoting expression of *Gad1, Gad2* (Le et al., 2017), *Nrxn3,* and *Arx* (Lindtner et al., 2019). Conversely, DLX2 represses expression of *Olig2*, a regulator for neural progenitor proliferation and oligodendroglial progenitor cell (OPC) differentiation (Petryniak et al., 2007). Although these studies support a role for DLX2 in regulating the neuronal *versus* glial fate determination during early neurogenesis, the mechanisms controlling gliogenesis remain unclear.

Meanwhile, Notch signalling plays an important role in balancing neurogenesis and gliogenesis (Grandbarbe et al., 2003). Activation of Notch signalling induces transcription of downstream TFs, such as *Hes1* and *Hes5* (Chillakuri et al., 2012), which in turn inhibit neurogenesis through repressing proneural factors (*Ascl1, Ngn2*). Conversely, Notch activation promotes cell proliferation and glial differentiation, thereby regulating neural progenitor fates (Grandbarbe et al., 2003; Shimojo et al., 2008).

Here, we show that DLX2 inhibits glial differentiation cues and directly impacts the neuronal-glial switch. Using a comprehensive multi-omics approach we identify that Notch signalling genes are directly repressed by DLX2 in the developing GE during neurogenesis. Spatial transcriptomics of E12.5-E14.5 wildtype (WT) and *Dlx1/Dlx2^-/-^* forebrain identified regionally specific Notch signalling dysregulation upon loss of *Dlx1/Dlx2*. Analyses of the transcriptome and chromatin accessibility at single nuclei resolution revealed enhanced glial gene signatures and glial-specifying TF activities in *Dlx1/Dlx2^-/-^* GE. DLX2 repression of Notch signalling and glial-promoting TFs was specific to neural progenitors and post-mitotic intermediate progenitors and is necessary for inhibiting glial cell fate. Thus, DLX2 functions to exert regulatory control on cell fate determination through a complex orchestration of Notch signalling and TF regulation in specific cell populations, thereby ensuring the correct temporal neural developmental sequence.

## Results

### Single cell spatial transcriptomic analyses identify regional specification and novel secondary proliferative communities in the developing telencephalon

We applied single cell whole genome spatial transcriptomics to assess alterations to spatiotemporal gene expression patterns in the developing mouse forebrain and GE lamination upon loss of *Dlx1/Dlx2* at E12.5-E14.5. The ability to preserve the tissue location of sequencing data provides a powerful approach to examine transcriptomic profiles of cell populations within their spatial context (Chen et al., 2022). Evaluation of canonical forebrain markers reflects established expression patterns in the WT telencephalon (**Figs. 1A-1B**). The progenitor marker *Fabp7* exhibited localised expression in the ventricular zone (VZ), with *Ascl1* and *Pax6* further delineating the GE and neocortex, respectively. Intermediate progenitor (IP) marker *St18* localised mostly to subventricular zone (SVZ) in the GE, while dorsal IP (dIP) marker *Eomes* expression was specific to neocortical SVZ (**Fig. 1B**). The immature neuron marker *Dcx* localised to the MZ, while the dorsal neuronal marker *Tbr1* were expressed exclusively in the neocortical MZ (**Fig. 1B**). *Olig2* and *Nkx2-1*, two ventral telencephalon markers, were localised to the MGE (**Fig. 1B**). *Dlx1/Dlx2* expression was localised to the GE across the E12.5-E14.5 embryonic time points in WT; expression was not detected in *Dlx1/Dlx2^-/-^* tissue (**Fig. S1A**). In *Dlx1/Dlx2^-/^*^-^ tissue, the expression patterns of marker genes mirrored that of the WT tissue, with increased expression of *Olig2* and *Nkx2-1* (**Fig. 1B**). To identify cell types, unsupervised clustering and manual annotation were performed based on cell type markers, defining 11 distinct clusters where the distribution of each cell type corresponded to the anatomical regions that have been extensively described(Bandler et al., 2017; Jiang & Nardelli, 2016). NPs and IPs were localised in the VZ and SVZ, respectively, and post-mitotic neurons were found in the mantle zone (MZ) (**Fig. 1C**). Visualisation of cell type clusters from WT and *Dlx1/Dlx2^-/-^* forebrains showed significant separation consistent with the defined anatomic locations (**Fig. 1D**). In the ventral telencephalon, two ventral IP (vIP) populations were identified, segregated by their proliferative state shown by differences in *Mki67* and *Top2a* expression (**Figs. S1B-S1C**). These data significantly elaborate on previously established forebrain models through our identification of a bespoke secondary proliferative layer within the ventral SVZ (**Figs. 1C-1E**). Decreasing expression levels of progenitor markers (*Fabp7*, *Nes*, *Ascl1*) and increasing expression of IP and neuronal markers (*St18*, *Dcx*) across ventral NP (vNP), vIP2, and vIP1 further support the transitional cell states of vNP, vIP2, and vIP1 (**Fig. 1E**). The dorsal neural progenitors (dNP), dIP, and dorsal neurons (dNeurons) were marked by *Pax6*, *Eomes*, and *Tbr1*, respectively (**Fig. 1E**). Although OPC (*Pdgfra*, *Sox10*) and astroglial marker (*Sox9, Nfia*) expression were detected, no distinct glial cells were identified in either WT or *Dlx1/Dlx2^-/-^* tissue (**Fig. S1B**).

**Figure 1.**
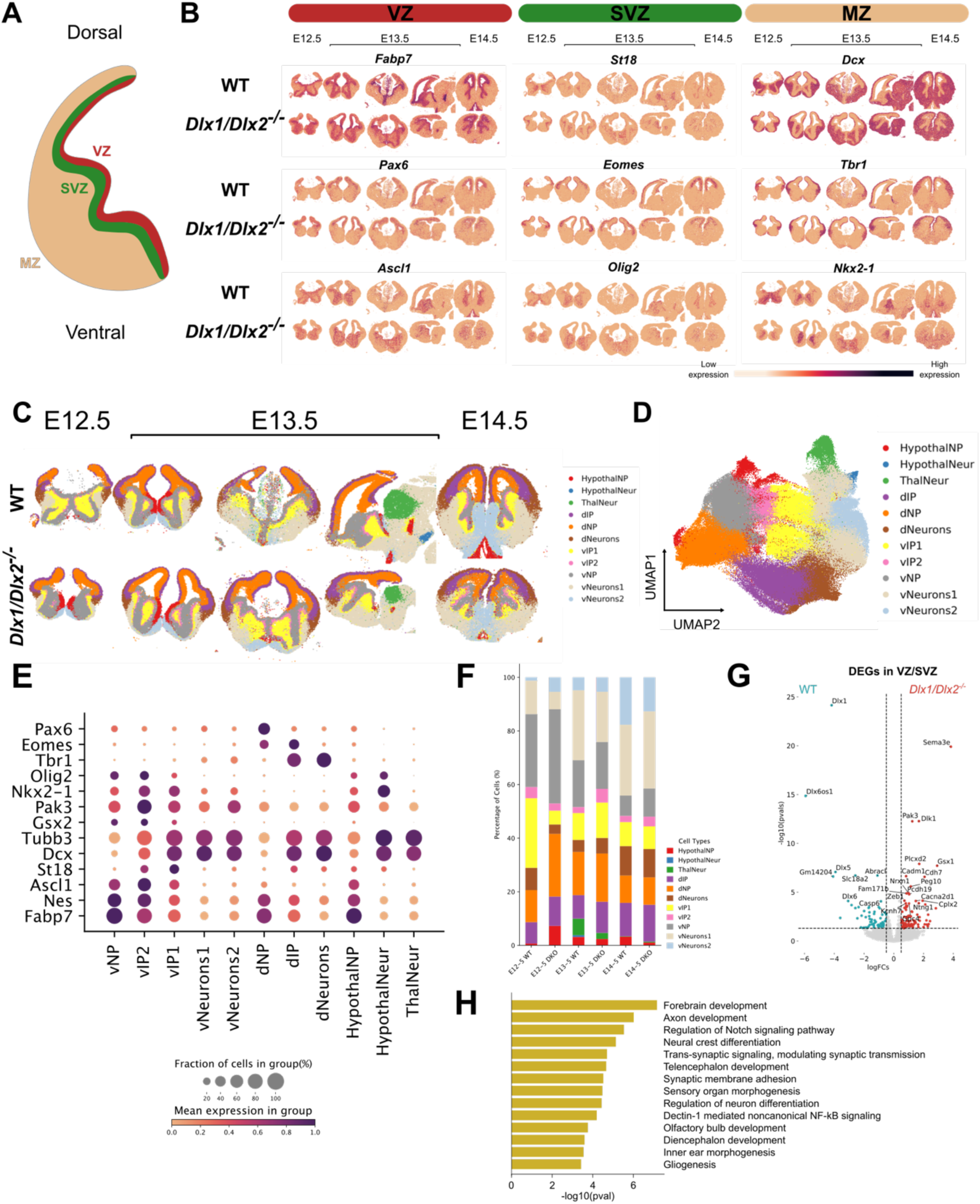
Spatiotemporal resolution in the developing forebrain. **(A)** Schematic diagram illustrating the sub-anatomical compartments within the embryonic telencephalon, including the ventricular zone (VZ), sub-ventricular zone (SVZ), and mantle zone (MZ). **(B)** Expression of sub-anatomical markers across sections from E12.5 (left), E13.5 2x coronal, 1x sagittal (middle), E14.5 (right). From left to right column: VZ markers, SVZ markers, and MZ markers. From top to bottom row: telencephalon markers (VZ: *Fabp7*, SVZ: *St18*, MZ: *Dcx*); dorsal telencephalon markers (VZ: *Pax6*, SVZ: *Eomes*, MZ: *Tbr1*), and ventral telencephalon markers (VZ/SVZ: *Ascl1, Olig2*, MZ: *Nkx2-1*). **(C)** Cell type classification following unsupervised clustering and manual annotation. WT sections (top row) and *Dlx1/Dlx2^-/-^* sections (bottom row) for E12.5 (left), E13.5 (2x coronal, 1x sagittal) (middle) and E14.5 (right). v/dNP: ventral/dorsal neural progenitors, v/dIP: ventral/dorsal intermediate progenitors, ThalNeur: thalamic neuron, HypothalNeur: hypothalamic neuron. **(D)** UMAP plots showing clustering and cell type classification for WT and *Dlx1/Dlx2^-/-^* sections. Legend as shown in (C). **(E)** Dot plot showing the mean expression of cluster markers in cells in each cluster from (C). Dot sizes denote the fraction of cells expressing the corresponding markers. **(F)** Bar plot showing the percentage of each cell type in WT and *Dlx1/Dlx2^-/-^* E12.5-E14.5 spatial transcriptomics dataset, summarised based on age and genotype. Legend in (C). **(G)** Volcano plot showing the differential expression analyses comparing VZ and SVZ of WT vs *Dlx1/Dlx2^-/-^* GE. Thresholds for differentially expressed genes were set at FDR<0.05 and fold change > 1 or < -1. **(H)** Gene ontology analysis results of all differentially expressed genes in VZ and SVZ of *Dlx1/Dlx2^-/-^* GE.

Overall, the cell type proportions showed an increase of post-mitotic neurons and a decrease of neural progenitors from E12.5-E14.5. In *Dlx1/Dlx2^-/-^* GE there was a decrease in the number of vNeurons, as *Dlx1/Dlx2* promotes GABAergic IN differentiation (**Fig. 1F**). There was also a subtle increase in vNPs and vIPs in E13.5 and E14.5 *Dlx1/Dlx2^-/-^* GE, whereas E12.5 *Dlx1/Dlx2^-/-^*GE displayed fewer vIPs. However, changes in the E12.5 GE may reflect differences in tissue sections rather than biological differences. Despite these changes in cell proportions, the trilaminar organisation of the *Dlx1/Dlx2^-/-^*GE did not display significant changes compared to WT. To identify underlying molecular alterations underpinning the increase in vNPs and vIPs in *Dlx1/Dlx2^-/-^*GE, gene expression was compared in the VZ and SVZ of the WT and *Dlx1/Dlx2^-/-^*GE (**Fig. 1G**). Differentially expressed genes (DEGs) in *Dlx1/Dlx2^-/-^* VZ and SVZ were significantly enriched in forebrain development and axon development as well as Notch signalling pathway genes, which included the upregulation of *Dlk1, Dll4, Jag1,* and *Hes6* (**Figs. 1H, S2; Table S1**). Enhanced Notch signalling reflects increased cell proliferation and reduced neuronal differentiation, thereby suggesting its potential role in inhibiting vIP differentiation, contributing to the observed decrease numbers of neurons. Meanwhile, enrichment of DEGs implicated in gliogenesis, neural crest differentiation, and development of other neural structures such as olfactory bulb and diencephalon suggests an altered progenitor cell fate in the *Dlx1/Dlx2^-/-^* forebrain (**Fig. 1H; Table S1**). These findings substantiate the role for *Dlx1/Dlx2* in ensuring proper spatial expression of neural differentiation genes in the VZ and SVZ (Anderson, Qiu, et al., 1997).

### Identification of cell communities that form distinct regional relationships

To further understand the spatial organisation between specific cell populations, we examined regions of the developing brain composed of sub-anatomical cell communities(Farah et al., 2024). Cell community analysis was performed by clustering tissue areas based on the proportion of cell types in each area, resulting in 15 cell communities across all sections and time-points (**Fig. 2A**). Some cell communities consisted primarily of one cell type, e.g. communities 8 (thalamic neurons (ThalNeur)) and 9 (vIP1), whereas other cell communities propose a refined classification of sub-anatomical regions, further dividing the VZ-SVZ-MZ layers of the developing telencephalon. In the GE, communities 0, 7, and 11 contained different proportions of vNPs, vIPs, and vNeurons (**Figs. 2A-2B**). Community 0 was mostly comprised of vIP1s (56%) with a low proportion of vNPs, vIP2s and vNeurons, whereas community 7 mostly contained vNPs (68%), and community 11 was primarily vIP2s (65%) with a secondary proportion of vNPs (29%) (**Table S2**). The differences in cell type proportions between communities were also reflected by their location, where community 7 was closest to the VZ and community 0 was closest to the MZ (**Fig. 2A**). Such analysis demonstrates the presence of tissue sub-layering underneath the VZ-SVZ-MZ axis, and the cell communities identified illustrate the heterogeneity and complex organisation of the developing forebrain.

**Figure 2.**
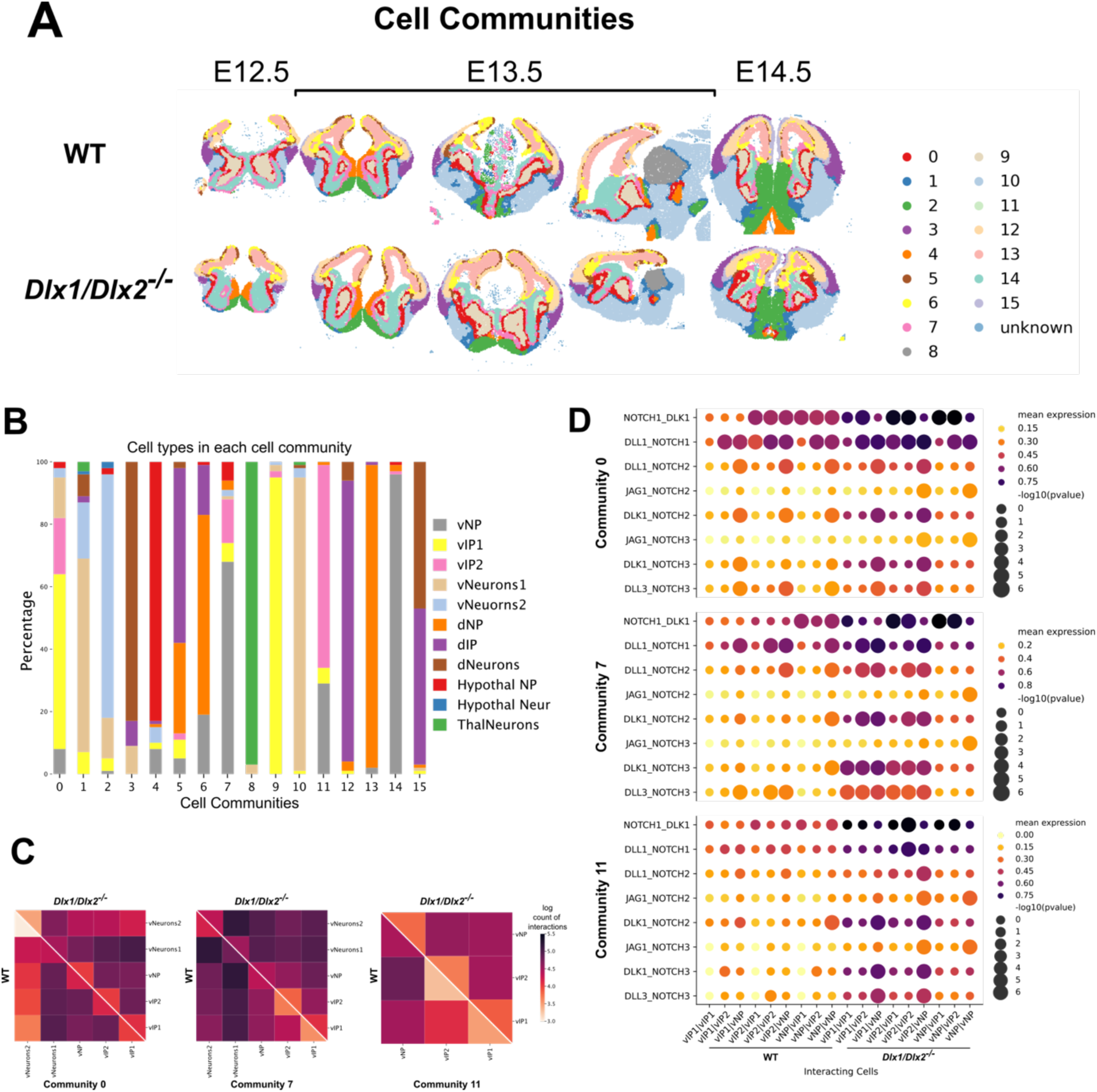
Novel cell communities and spatially dysregulated Notch signalling in the absence of *Dlx1/Dlx2* function. **(A)** Clustering results from cell community analysis, identifying 15 distinct clusters and one unknown population. Top row: WT, bottom row: *Dlx1/Dlx2^-/-^* sections. **(B)** Proportion of each cell type in each of the 15 cell communities identified. **(C)** Heatmap showing log count of interactions between vNPs, vIP1, vIP2, vNeurons1, and vNeurons2 in communities 0, 7, and 11. Bottom half of each heatmap represents the interactions in WT GEs and top half represents the interactions in *Dlx1/Dlx2^-/-^* GE. **(D)** Dot plot showing Notch ligand-receptor interaction between vNPs, vIP1, and vIP2 in communities 0, 7, and 11. Color of each dot represents the mean expression of the ligand and receptor between interacting cell types, whereas dot size represents the statistical significance of interaction (-log_10_(pvalue)). The order of interacting cell type label (x-axis) corresponds to the cell type that expressed the molecule in the order of the ligand-receptor label (y-axis).

### Sufficient Notch signalling levels are required for the differentiation of proliferating cell populations in the ganglionic eminences

We focused on the cell communities adjacent to the SVZ of the GE, communities 0, 7, and 11, which consisted of a mixture of vIPs and vNPs, to better characterise Notch signalling dysregulation in *Dlx1/Dlx2^-/-^* telencephalon. We found that across these communities the intra-cell type interaction was increased in the *Dlx1/Dlx2^-/-^* GEs, notably between vIPs and vNPs in community 7, and between vIPs in community 11. In community 0, there were increased interactions between vNeurons and other cell types, the most apparent being the interactions between vNeurons and vIP1 (**Fig. 2C**). SHH, WNT, BMP, Notch, and Semaphorin ligand-receptor interactions were identified across all three GE communities (**Table S3**), all common signalling pathways present in the developing telencephalon.

Given the upregulation in Notch signalling in VZ and SVZ in *Dlx1/Dlx2^-/-^*GE (see Fig. 1G), we explored the spatial context of Notch dysregulation in the GE communities. In WT GEs, Notch signalling was enriched in communities 0 and 7 between vIP and vNP via NOTCH1, activated by DLL1 or DLK1 (**Fig. 2D**). In *Dlx1/Dlx2^-/-^*GE, there was an increase of Notch signalling by vIP2 to vNP across all communities. In community 7, most proximal to the VZ, vNPs and vIP2 appeared to receive stronger signalling via DLK1 and DLL1 from vIP1 and vIP2. Furthermore, there was a gain of canonical Notch signalling between vIP1 and vIP2 in *Dlx1/Dlx2^-/-^* GE through DLL1/DLL3 and NOTCH1/NOTCH2/NOTCH3 interactions and vIP1-vIP1 interactions through signals received by NOTCH3. In community 0, closest to the MZ, signalling received by *Dlx1/Dlx2^-/-^* vIP2 and vNP via DLL1-NOTCH1 interactions was the most upregulated (**Fig. 2D**). Differential Notch ligand-receptor interactions in community 11 of *Dlx1/Dlx2^-/-^* GE were highly specific, where signalling via NOTCH1 only increased between vIP2-vIP2, while signalling via NOTCH2/NOTCH3 was elevated in vIP1-vNP and vIP2-vNP interactions. Noticeably, JAG1 signalling increased in vNP-vNP interactions of all three communities via interacting with NOTCH2/NOTCH3 receptors. These findings support the upregulation of Notch signalling in *Dlx1/Dlx2^-/-^* GE via differentiated IPs (vIP1) and was primarily received by the proliferating IPs (vIP2) and vNP. Most increased interactions were also observed in regions closer to the VZ, where a greater number of proliferating cells (vNP and vIP2) were found. Since Notch signalling promotes cell proliferation through lateral inhibition of differentiation, this likely contributed to the increased number of *Dlx1/Dlx2^-/-^*vNPs and vIPs. This Notch ligand-receptor analysis from cell communities 0, 7, and 11 showed the upregulation of Notch in the absence of *Dlx1/Dlx2* has spatially specific effects that directly impact neuronal cell fate. The proliferative signals from the increased Notch signalling promote maintenance of NPs closer to the VZ, whereas in SVZ areas more proximal to the MZ, these signals resulted in an arrest of vIP differentiation. Collectively, these findings illustrate the contextual spatial significance in neuronal fate regulation and allow us to define specific populations and their regional identity.

### Dysregulated Notch signalling in *Dlx1/Dlx2^-/-^* GE impairs neural cell fate specification

To further understand the impact of increased Notch signalling upon loss of *Dlx1/Dlx2* in the GE, differential gene networks and differentiation trajectories were mapped at single nuclei resolution across E12.4 to E14.5 in WT and *Dlx1/Dlx2^-/-^* GE (**Figs. 3A-3B, S3A**). After quality control filtering, we obtained 18,592 cells from WT and 14,546 cells from *Dlx1/Dlx2^-/-^*GEs. Across three embryonic timepoints, 7,204 cells were analysed from E12.5, 16,491 cells from E13.5, and 9,443 cells from E14.5 GEs (**Fig. 3B (i)**). The UMAP of WT and *Dlx1/Dlx2^-/-^* samples occupied similar space showing cell types present in both data sets were closely related (**Fig. 3B (ii)**). Larger proportions of cells in G2/M and S phase were found at E12.5, whereas more cells in G1 phase were found at E13.5 and E14.5 reflecting greater numbers of cycling progenitors at the earlier timepoint (**Fig. 3B (iii)**). Unsupervised clustering followed by manual annotation using expression of known markers(Bandler et al., 2017; Jiang & Nardelli, 2016) identified 9 cell types across all samples (**Fig. 3C**). Most post-mitotic cells from the GE were GABAergic INs expressing *Gad1, Gad2,* and *Nrxn3*, with a separate population also expressing *Sst* and *Npy* (*Sst+ Npy+* INs) (**Figs. 3C-3F**). An IN population lacking *Gad* expression was also identified (*Gad-* INs). Two separate ventral progenitor populations, radial glia and ventral neural progenitors (NPs), were identified, distinguished by their cell cycling phases. The radial glia population corresponded to cells in G2/M phase and ventral NP cells to those in S phase (**Figs. 3C-3F**). A population of early differentiated ventral intermediate progenitors (IP) was also identified, as marked by expression of *Dcx* and *Nkx2-1* (**Figs. 3C-3F**). The only post-mitotic glial population identified comprised of oligodendroglial precursor cells (OPCs), marked by expression of *Olig2, Pdgfra, Sox10*, and *Cspg4* (**Figs. 3D-3F**). Populations of dorsal cell types were also present, with dorsal IPs marked by expression of *Eomes* and glutamatergic neurons marked by *Tbr1* (**Figs. 3D-3F**).

**Figure 3.**
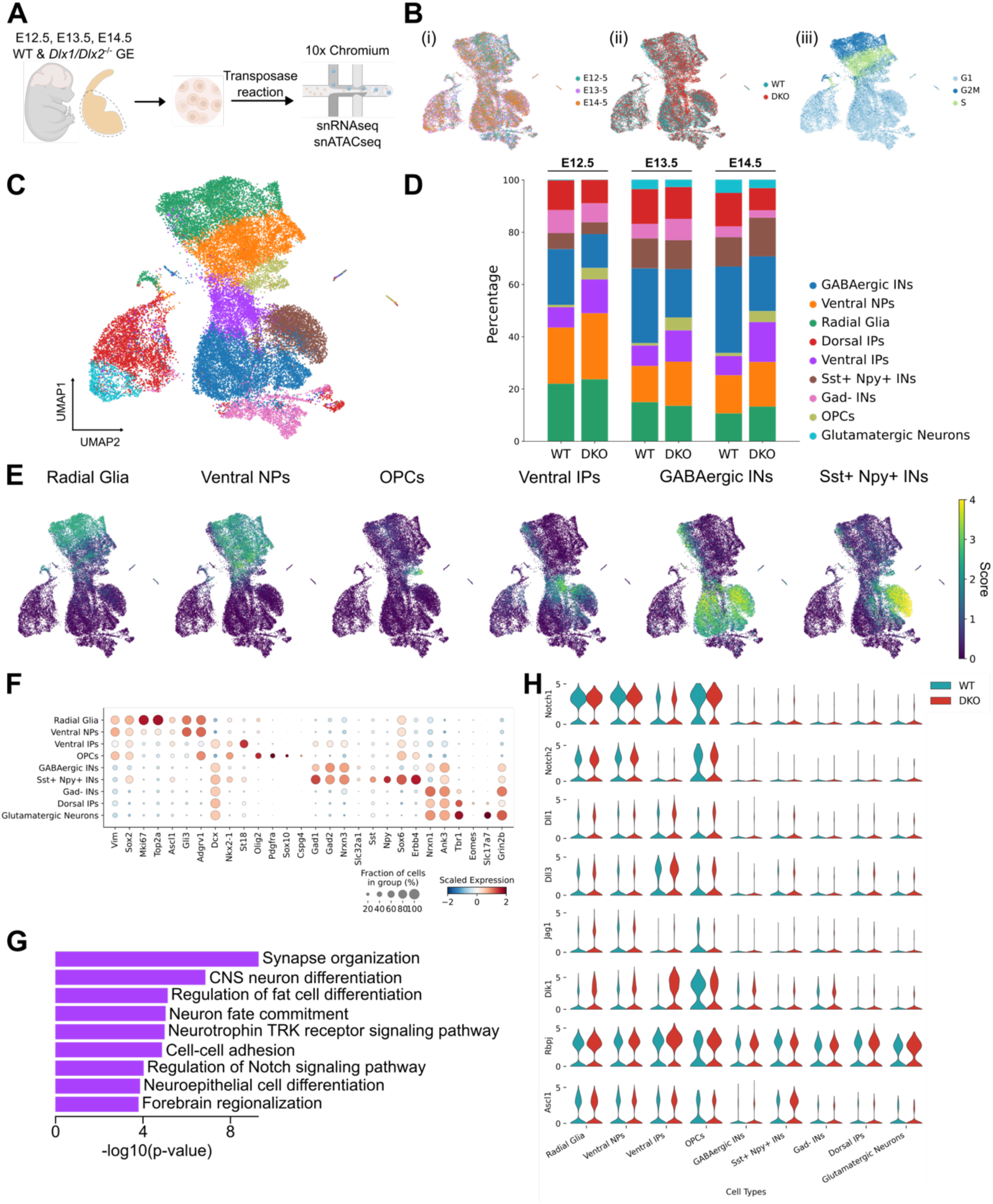
Increased Notch signalling and altered neuronal-glial cell fate in *Dlx1/Dlx2^-/-^* embryonic mouse GE. **(A)** Schematic of experimental design for snMultiome. GEs from E12.5-E14.5 *Dlx1/Dlx2^-/-^* and WT littermate controls were assessed. E12.5 and E14.5 (n=1); E13.5 (n=2). **(B)** UMAPs of E12.5-E14.5 snMultiome dataset, cells colored by (i) embryonic age, (ii) genotype, and (iii) cell-cycle state. **(C)** UMAP of E12.5-E14.5 snMultiome dataset, cells colored by cell type. Legend in (D). **(D)** Bar plot showing the percentage of each cell type in WT and *Dlx1/Dlx2^-/-^* E12.5-E14.5 snMultiome dataset. NPs: neural progenitors, IPs: intermediate progenitors, OPCs: oligodendroglial precursor cells, IN: interneurons. **(E)** Visualisation of cell type signature scores for radial glia (*Vim, Sox2, Mki67, Top2a*), ventral NPs (*Ascl1, Gli3, Adgrv1*), OPCs (*Olig2, Sox10, Pdgfra, Cspg4*), ventral IPs (*Dcx, Nkx2-1, St18*), GABAergic INs (*Gad1, Gad2, Slc32a1, Nrxn3*), and *Sst+ Npy+* INs (*Sst, Npy, Sox6, Erbb4*). **(F)** Dot plot showing the expression of cell type marker genes. **(G)** GO analysis for upregulated genes in *Dlx1/Dlx2^-/-^* ventral IPs. **(H)** Violin plots showing expression of Notch signalling pathway genes in E12.5-E14.5 WT and *Dlx1/Dlx2*^-/-^ GEs. WT: wildtype, DKO: *Dlx1/Dlx2^-/-^*.

Loss of *Dlx1* and *Dlx2* expression was confirmed in *Dlx1/Dlx2^-/-^*GEs (**Fig. S3A**), along with an expected reduction in the number of GABAergic INs (**Fig. 3D**) (Le et al., 2017). Notably, the number of *Sst+ Npy+* GABAergic INs was not affected in the mutants despite the downregulation of most *Sst+ Npy+* IN specification TFs (*Lhx6, Lhx8, Sox6, Runx1tl*), likely due to the upregulation of *Nkx2-1*, another key TF for *Sst+ Npy+* INs specification (**Figs. 3D, S3B**). Similar to changes observed in our spatial transcriptomics dataset, radial glia, ventral NPs, and ventral IPs proportions were increased in the *Dlx1/Dlx2^-/-^* GE (**Figs. 1F, 3D**). In addition, the number of OPCs also increased in *Dlx1/Dlx2^-/-^* GE, supporting upregulated gliogenesis in the spatial transcriptomics DEGs (**Figs. 1G, 3D**). Dysregulation in the expression of many neuronal genes in ventral progenitors and ventral IPs expression was seen, suggesting disrupted neuronal differentiation processes (**Fig. S3C; Table S4**). Again, consistent with our spatial transcriptomics findings, upregulated DEGs of ventral progenitors and IPs enriched for genes implicated in neuronal differentiation and fate commitment, including Notch signalling genes such as *Dll1, Dll3, Dlk1, Jag1,* and *Hes6,* which were also upregulated in *Dlx1/Dlx2^-/-^* OPCs (**Figs. 3G-3H; Table S4**). Conversely, Notch signalling genes were expressed minimally in INs (**Fig. 3H**). In ventral progenitors and IPs, there was also an enrichment of TFs important for diencephalon development amongst the upregulated DEGs, including *Gsx1, Isl1* and *Otp*, indicating that lack of *Dlx1/Dlx2* also affects neuronal fate specification (**Figs. S3D-S3E; Table S4**). Conversely, axonogenesis and GABAergic neuron differentiation genes were enriched in the downregulated DEGs of the *Dlx1/Dlx2^-/-^* progenitors and ventral IPs (**Figs. S3D-S3E; Table S4**). DEG enrichment in ventral progenitors and IPs along with increased OPC populations reveal that neuronal and glial cell fate determination processes were impaired, correlated with significantly dysregulated Notch signalling pathways.

### *Dlx1/Dlx2* are necessary for promoting neuronal differentiation and concomitantly repressing glial fate in committed neural progenitors

Since changes in cell populations in *Dlx1/Dlx2^-/-^* GE suggest altered progenitor cell differentiation, differentiation trajectories of the ventrally derived WT and *Dlx1/Dlx2^-/-^* cell types were examined. Both WT and *Dlx1/Dlx2^-/-^* cells presented a general differentiation trajectory of radial glia into ventral NPs, ventral IPs, and finally GABAergic INs (**Fig. 4A-4B**). However, *Dlx1/Dlx2^-/-^*GE demonstrate a unique OPC differentiation branch and instead of a single branch for GABAergic INs evident in WT tissue, display three differentiation branches, with the third branch depicting the *Sst+ Npy+* IN population (**Fig 4A-4B**). The WT and *Dlx1/Dlx2^-/-^* progenitors were then projected onto the same differentiation path to enable a direct comparison of differentiation trajectories, separated by cell type and differentiation segment (**Figs. 4C-4D**). This analysis reveals that the OPCs and *Sst+ Npy+* INs branches predominantly consisted of cells from *Dlx1/Dlx2^-/-^* GE, demonstrating the altered differentiation trajectories of progenitor cells in the absence of *Dlx1/Dlx2* (**Fig. 4E**). Similarly, GABAergic INs were predominantly from WT GE. Furthermore, pseudotime scores along the differentiation tree showed that *Dlx1/Dlx2^-/-^* ventral IPs were less mature than WT ventral IPs (**Fig. 4F**) suggesting a stall in their differentiation. In contrast, GABAergic INs from WT and *Dlx1/Dlx2^-/-^* GE shared a similar pseudotime score indicating maturation of differentiated GABAergic INs were not affected by a lack of *Dlx1/Dlx2* function (**Fig. 4F**).

**Figure 4.**
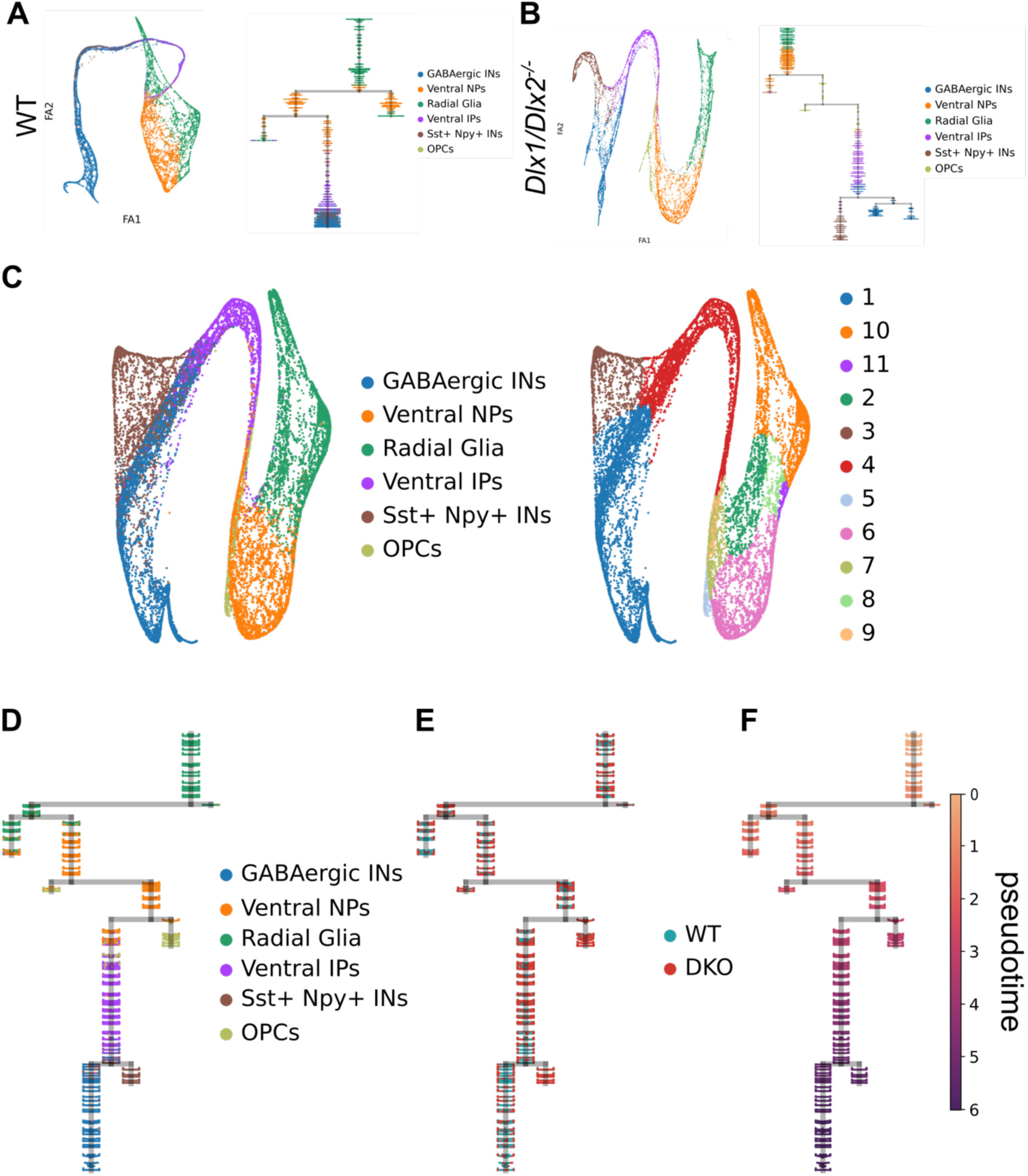
Altered neural progenitor cell differentiation trajectories upon loss of *Dlx1/Dlx2*. **(A)** Differentiation trajectories in WT GE represented with force-directed embedding (left), and differentiation dendrogram (right) showing differentiation of radial glia to ventral IPs, then GABAergic INs. INs: interneurons, NPs: neural progenitors, IPs: intermediate progenitors, OPCs: oligodendroglial precursor cells. **(B)** Differentiation trajectories in *Dlx1/Dlx2^-/-^*GE represented with force-directed embedding (left), and differentiation dendrogram (right) showing OPCs and *Sst+ Npy+* INs differentiation branches and GABAergic INs differentiation branches are split into two in comparison to a single branch in WT. **(C)** Force-directed embedding of the ventrally derived cell types in WT and *Dlx1/Dlx2^-/-^* GE used for differentiation trajectory analysis, showing clusters coloured by cell types (left) and differentiation segments (right). **(D)** Dendrogram depicting the differentiation trajectories of ventral forebrain cell types from E12.5-E14.5 WT and *Dlx1/Dlx2^-/-^*GE snRNAseq dataset colored by cell type. **(E)** Dendrogram depicting the differentiation trajectories of ventral forebrain cell types coloured by genotype. WT: wildtype, DKO: *Dlx1/Dlx2^-/-^*. **(F)** Dendrogram depicting the differentiation trajectories of ventral forebrain cell types coloured by pseudotime score.

Given the significant differences in the WT and *Dlx1/Dlx2^-/-^*ventral IP and OPC lineages, we next examined the expression of individual IN and glial differentiation regulators to further characterise changes to cell fates upon loss of *Dlx1/Dlx2*. *Dlx1/Dlx2^-/-^* ventral IPs showed altered expression of forebrain patterning TFs, such as increased *Isl1*, *Otp*, and *Nkx2-1* and reduced *Erbb4* and *Lhx6* expression, indicating reduced specification of MGE-derived INs, contributing to the reduced number of GABAergic INs (**Fig. 5A**). In *Dlx1/Dlx2^-/-^* OPCs, we observed downregulation of *Cspg4* (early OPC marker) and upregulation of *Sox10* (late OPC marker), as well as *Olig2* and *Pdgfra* which were also upregulated in radial glia and ventral NPs indicating an adoption of OPC fate in *Dlx1/Dlx2^-/-^* neural progenitor cells (**Fig. 5A**). Furthermore, astroglial markers including *Fabp7*, *Sox9*, and *Notch1* were increased in *Dlx1/Dlx2^-/-^* radial glia, ventral NPs, and ventral IPs, but the astrocyte lineage marker *Gfap* was not expressed in WT or *Dlx1/Dlx2^-/-^*GE (**Fig. 5A**). Upregulation of astroglial markers and increased OPCs during neurogenesis indicate early adoption of gliogenesis in *Dlx1/Dlx2^-/-^* GE, defining a role for DLX1/DLX2 in the concurrent regulation of neuronal and glial cell fates. Thus, we have identified cell populations with altered marker gene expression thereby revealing *Dlx1/Dlx2* temporal regulation of cell fate determination during neurogenesis.

**Figure 5.**
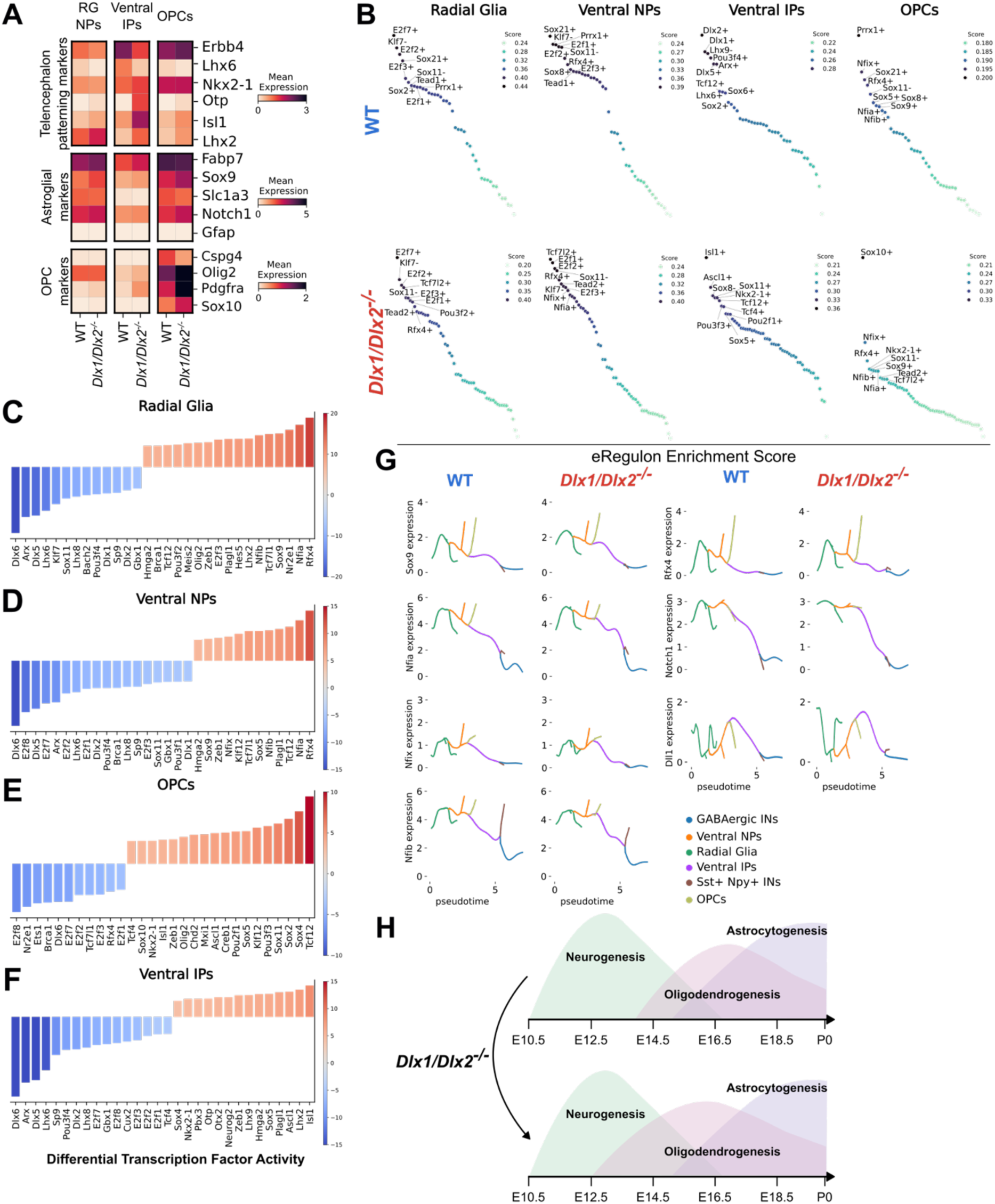
Altered neural cell fate in *Dlx1/Dlx2^-/-^* GE is driven by changes in eRegulon and TFs activities. **(A)** Heatmap showing mean expression of telencephalon patterning markers (top: *Erbb4, Lhx6, Nkx2-1, Otp, Isl1, Lhx2*), astroglial markers (middle: *Fabp7, Sox9, Slc1a3, Notch1, Gfap*) and OPC markers (bottom: *Cspg4, Olig2, Pdgfra, Sox10*); and in radial glia and ventral NPs (left), ventral IPs (middle), OPCs (right). **(B)** Ranked eRegulon specificity score for radial glia, ventral NPs, ventral IPs, and OPCs in E12.5-E14.5 WT and *Dlx1/Dlx2^-/-^* GE. Positive or negative gene regulation denoted by ‘+’ or ‘-’. **(C-F)** Bar plots showing differential TF activity in radial glia (C), ventral NPs (D), OPCs (E), and ventral IPs (F) in E12.5-E14.5 *Dlx1/Dlx2^-/-^* (red) vs WT (blue) cells. **(G)** Expression of *Sox9, Nfia*, *Nfix, Nfib, Rfx4, Notch1*, and *Dll1* in E12.5-E14.5 WT and *Dlx1/Dlx2^-/-^* GE along the ventral forebrain cell differentiation trajectories (see Fig. 4D-4F). **(H)** Schematic of the temporal neural development sequence, where neurogenesis starts at mid-gestation (E10.5), followed by the beginning of oligodendrogenesis at E14-E14.5 and astrocytogenesis at E15.5-E16. Deletion of *Dlx1/Dlx2* results in oligodendrogenesis starting as early as E12.5.

### Altered enhancer-driven gene regulatory network activities in *Dlx1/Dlx2^-/-^* GE directly impact neural cell fate decisions

Throughout development, cell-type specific gene expression is often determined by enhancer occupancy through a combinatorial TF code (Catta-Preta et al., 2025; Nord & West, 2020). Hence, we took advantage of the coupled snRNA-snATAC-seq data to examine enhancer-driven gene regulatory networks (eRegulons) using SCENIC+ (Bravo Gonzalez-Blas et al., 2023) to identify TFs that directly impacted neural cell fate upon loss of *Dlx1/Dlx2*. WT radial glia and ventral NPs showed similar eRegulon enrichment of E2F TFs, SOX21, and SOX11 eRegulons (**Fig. 5B**). The E2F TFs direct cell cycle exit and promote egress from quiescent states in adult neural stem cells (Fong et al., 2022). Similarly, SOX21 and SOX11 play a role in neural progenitor maintenance (Mu et al., 2012). Enrichment for E2F family eRegulons in *Dlx1/Dlx2^-/-^* radial glia and ventral NPs were similar to WT, whereas the TCF7L2 eRegulon was highly enriched in both cell types only upon loss of *Dlx1/Dlx2* (**Fig. 5B**). In WT ventral IPs, several homeobox eRegulons were enriched, including DLX TFs, LHX6, and SOX6 (**Fig. 5B**), all regulators of *Sst+ Npy+* INs specification (Munguba et al., 2023). In contrast, eRegulons enriched in the *Dlx1/Dlx2^-/-^* ventral IPs differed from WT counterparts, including but not limited to loss of DLX eRegulons, and gain of alternative GABAergic IN promoting eRegulons, including NKX2-1, ISL1, and ASCL1 (**Fig. 5B**).

OPCs were enriched for glial cell differentiation regulators, including NFIA, NFIX, and SOX9 eRegulons. Interestingly, SOX10 and NKX2-1 eRegulons were exclusively found in the *Dlx1/Dlx2^-/-^* and not WT OPCs (**Fig. 5B**). The TCF7L2 eRegulon, found only in *Dlx1/Dlx2^-/-^* radial glia and ventral NPs, was also enriched in both WT and *Dlx1/Dlx2^-/-^* OPCs, underscoring its role in glial differentiation. The NFI TFs eRegulons, including NFIA and NFIX, function similarly in glial maturation and were also enriched in the *Dlx1/Dlx2^-/-^* but not WT ventral NPs, showing that they may be contributing to enhanced OPC maturation (**Fig. 5B**). These differential eRegulon enrichment showed that the altered cell fate in the absence of *Dlx1/Dlx2* is partially driven by changes in transcriptional programming.

Differential TF activities in WT vs *Dlx1/Dlx2^-/-^* GE were quantified to further understand the impact of each eRegulon on cell fate specification (**Figs. 5C-5F**). Notably, NFIA and RFX4 were TFs with the most upregulated activities in the *Dlx1/Dlx2^-/-^* ventral NPs and radial glia, along with other glial fate promoting TFs such as NFIX, SOX9, TCF7L1, and TCF12 (**Figs. 5C-5D**). Although the activities of these TFs were all upregulated in radial glia and ventral NPs, these TFs displayed different transcriptional changes in each cell type (**Figs. S4A-S4B**). Activities of established factors that promote OPC differentiation and maturation (OLIG2, SOX10, TCF12) and migration (SOX5) also displayed upregulated activities in the *Dlx1/Dlx2^-/-^* OPCs, with largely unique gene targets among these TFs (Baroti et al., 2016) (**Figs. 5E, S4C**). These changes show that the increased number of OPCs in *Dlx1/Dlx2^-/-^* GE were driven by an overall elevated OPC differentiation programme in both ventral progenitors and OPCs. Consistent with the differential expression analysis (**Fig. S3B**), in *Dlx1/Dlx2^-/-^* ventral IPs activities of *Sst+ Npy+* INs regulators LHX6 and LHX8 decreased; meanwhile SOX6 activity was maintained, and certain dorsal (LHX2, NEUROG2) and diencephalon specific TFs (ISL1, OTP) had increased activities (**Figs. 5F, S4D**). These data illustrate how *Dlx1/Dlx2* impacts cell fate decisions by ensuring tight regulation of transcriptional programs in the developing GE.

### The neuronal-glial fate switch is promoted prematurely in *Dlx1/Dlx2^-/-^* GE

During neural development, neurogenesis typically occurs prior to gliogenesis, including both oligodendrogenesis and astrocytogenesis. Our differentiation trajectory (**Figs. 4D-4F**) and TF eRegulon analyses (**Figs. 5B-5F**) revealed that neuronal differentiation was stalled in ventral IPs and that glial differentiation programs were more active in progenitors in the absence of *Dlx1/Dlx2*. These data suggest that neurogenesis processes were halted and gliogenesis was initiated early in the *Dlx1/Dlx2^-/-^* GE. Additionally, we have identified that DLX2 regulates cell fate determination via repression of the Notch signalling pathway. We next examined expression of glial fate promoting TFs (*Sox9, Nfia, Nfib, Nfix*, *Rfx4*) and Notch signalling genes (*Notch1*, *Dll1*) along the differentiation trajectory previously constructed to better understand the dysregulated neuronal-glial timing in *Dlx1/Dlx2^-/-^* GE (**Figs. 4D, 5G**). Expression of each of these genes showed a similar temporal trend in WT GE, where expression increased during mid-radial glia, ventral NPs, OPCs, and late ventral IPs. Similar trends were observed in the *Dlx1/Dlx2^-/-^* GEs except increased expression of *Sox9, Nfia, Notch1*, and *Dll1* in the ventral IPs was shifted relative to WT to an earlier trajectory (**Fig. 5G**). *Nfib, Nfix*, and *Rfx4* displayed similar temporal changes in their expression patterns (**Fig. 5G**). The shift in the expression of glial fate promoting TFs along with its enhanced activities in the *Dlx1/Dlx2^-/-^* GE supports disruption of this temporal sequence, with premature glial differentiation in the GE (**Fig. 5H**).

### DLX2 controls the neuronal-glial fate switch through direct repression of Notch signalling pathway and glial promoting TFs

To identify DLX2 downstream targets in controlling neuronal-glial fate decisions, we performed DLX2 CUT&RUN, alongside H3K27ac ChIPseq, bulk RNAseq and ATACseq using E13.5 mouse GE (**Fig. 6A**). A consensus peak set containing 11,425 peaks from triplicate samples was merged to include all possible regions occupied by DLX2 (**Fig. S5A-S5C**). The distribution of the consensus peaks showed 36.11% at gene promoters (<3kb from transcription start sites (TSS)), 35.46% at distal intergenic regions, and 40.38% at intronic regions (**Fig. 6B, S5D**). *De novo* motif enrichment analysis showed that the DLX2 binding motif (TAATTA) was clearly the most significantly enriched in the consensus peak set, verifying the specificity of the assay (**Fig. 6C**). Verification of known DLX2 known targets including *Nrp2* and *Gad2* confirmed the specificity of the DLX2 CUT&RUN consensus peaks (**Fig. S5E**) (Le et al., 2007; Le et al., 2017).

**Figure 6.**
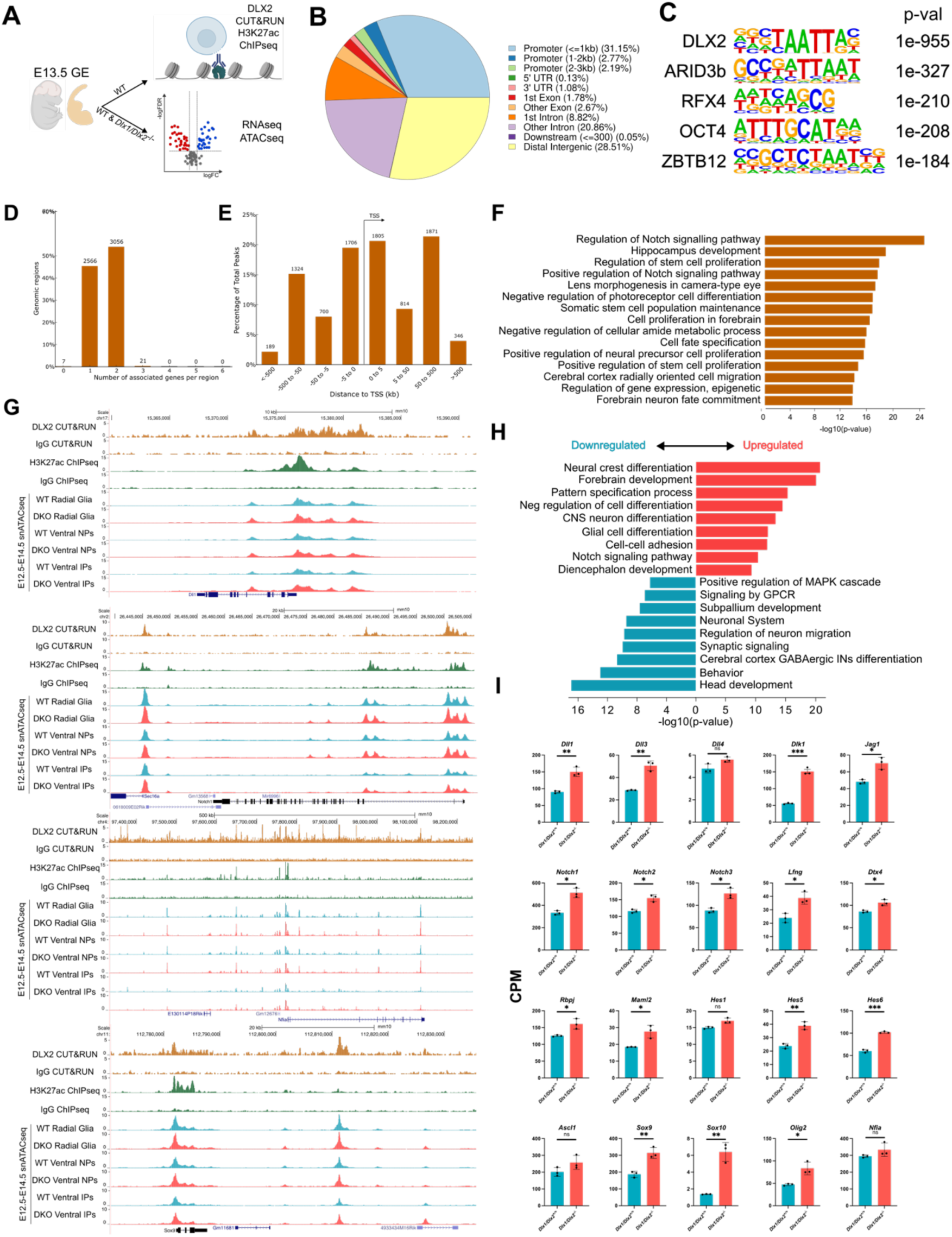
Identification of direct repression of Notch signalling and glial programs by DLX2 in the developing mouse GE. **(A)** Schematic of experimental design for DLX2 CUT&RUN, H3K27ac ChIPseq, RNAseq and ATACseq. **(B)** Pie chart of the genomic annotations for DLX2 CUT&RUN peaks. **(C)** Top 5 motifs from *de novo* motif enrichment analysis of DLX2 CUT&RUN peaks (n=3) by p-value. **(D)** Bar graph indicating the number of genes associated with both DLX2 CUT&RUN and H3K27ac ChIPseq consensus peaks. **(E)** Bar graph showing the genomic location of DLX2-H3K27ac-ATAC consensus peaks in relation to transcription start sites (TSS). **(F)** GO results of the consensus peaks from DLX2 CUT&RUN, H3K27ac ChIPseq, and ATACseq of E13.5 WT GE. **(G)** Coverage of DLX2 CUT&RUN, H3K27ac ChIPseq, and snATACseq of E13.5 WT and *Dlx1/Dlx2^-/-^* GE at *Notch1*, *Dll1*, *Nfia*, and *Sox9* loci. **(H)** GO analysis of up- and down-regulated genes in *Dlx1/Dlx2^-/-^* E13.5 GE. **(I)** Expression of genes involved in Notch signalling pathway and gliogenesis measured by RNAseq (counts per million, CPM) in E13.5 WT and *Dlx1/Dlx2^-/-^* mouse GE (n=3). *fdr<0.05, **fdr<0.005, ***fdr<0.0005.

We compared our DLX2 CUT&RUN dataset to a DLX2 ChIPseq study(Lindtner et al., 2019), and found 5756 overlapping peaks, corresponding to 3599 genes (**Fig. S6A**). DLX2 CUT&RUN demonstrated a higher percentage of peaks within promoters compared to DLX2 ChIPseq, whereas distal intergenic regions and intronic regions were more abundant with DLX2 ChIPseq (**Figs. S6B-S6C**). Both data sets showed comparable gene set enrichment, including cell fate specification, forebrain neuron differentiation, and neural precursor cell proliferation. However, DLX2 CUT&RUN also demonstrated enrichment in genes related to oligodendrocyte differentiation, which were not present in the ChIPseq data, showing the CUT&RUN captured a broader range of gene targets (**Fig. S6D**).

To identify active promoters and enhancers that DLX2 interacts with during neurogenesis, DLX2 CUT&RUN peaks were combined with data generated from ATACseq and H3K27ac ChIPseq in E13.5 WT GE, yielding 5652 peaks of DLX2-bound active regulatory regions. Of these DLX2-ATAC-H3K27ac consensus peaks, 2566 peaks corresponded to a single gene, whereas 3056 and 21 peaks corresponded to two and three genes, respectively (**Fig. 6D**). 3511 peaks were found to be within ±5kb away from TSS and 3195 peaks were found to be ±50-500kb away from TSS showing comparable active promoters and enhancer regions regulated by DLX2 (**Fig. 6E**). Strikingly, analysis of gene ontology terms revealed highly significant enrichment for regions associated with Notch signalling genes in open DLX2-ATAC-H3K27ac consensus peaks (**Fig. 6F**), including Notch receptor (*Notch1*), Notch ligands (*Dll1*, *Dlk1*, *Jag1*) and downstream TFs (*Hes1, Hey1, Ascl1*) (**Fig. 6G, S7A-S7D; Table S5**). Furthermore, DLX2-ATAC-H3K27ac consensus peaks also enriched for genes related to cell fate specification, including glial fate promoting TFs *Sox9, Nfia, Nfib,* and *Nfix* (**Fig. 6G, S7E-S7F**). The DLX2 bound regions of Notch signalling genes and glial fate promoting TFs demonstrated minimal changes in chromatin accessibility in the *Dlx1/Dlx2^-/-^* radial glia, ventral NPs and IPs determined by snATACseq (**Figs. 6G, S7A-S7F**), and were not enriched in differentially accessible promoters identified via ATACseq of E13.5 WT and *Dlx1/Dlx2^-/-^*GE (**Figs. S7G-S7H**). *Dll1*-, *Notch1*- and *Ascl1*-promoter driven luciferase reporter activity *in vitro* was reduced in the presence of DLX2 confirming the repressive effect of DLX2 (**Fig. S7I**). These data demonstrate that these genes were differentially expressed due to direct repression of DLX2 rather than a change in chromatin accessibility. Bulk transcriptomics analyses further validated the changes observed in both snMultiome and single cell spatial transcriptomics, where neuronal genes were downregulated upon loss of *Dlx1/Dlx2*, and Notch signalling genes and glial cell differentiation TFs were upregulated (**Figs. 6H-6I**). Overall, these data demonstrate that DLX2 ensures the tight regulation of neuronal-glial specification timing in the GE during E12.5-E14.5 through directly repressing expression of glial promoting TFs and Notch signalling genes in ventral progenitors and IPs, concomitantly promoting GABAergic INs differentiation and maturation. Coupling the gene regulatory networks determined through snMultiome and spatial transcriptomics data further highlights the impacted glial promoting TFs in neural progenitors located in the VZ (**Fig. 7A**). These data capture the complex DLX2 downstream network and redefine the mechanism by which DLX2 controls spatiotemporal cell fate specification in the developing telencephalon.

**Figure 7.**
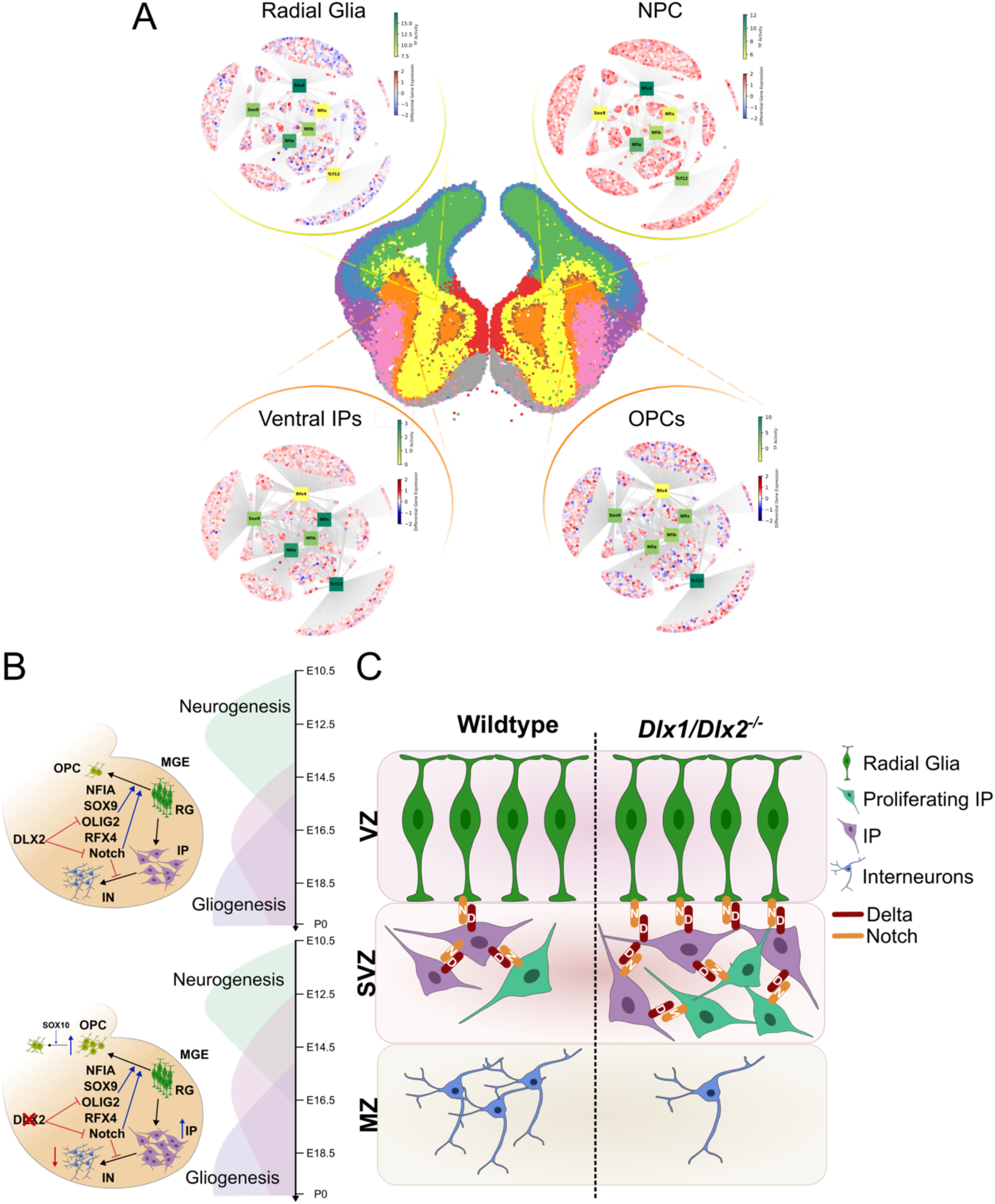
Model of DLX2 regulation of cell fate determination in the developing telencephalon at E12.5-E14.5. **(A)** Gene regulatory networks of selected glial differentiation promoting TFs (SOX9, NFIA, NFIB, NFIX, TCF12) exhibit varying activities in progenitors and IPs residing in the VZ and SVZ of the developing telencephalon. Specifically, these gene regulatory networks had enhanced activities in the neural progenitor cell (NPC) population in the VZ of the GE. **(B)** DLX2 promotes neuronal and represses glial differentiation through inhibiting Notch signalling genes and eRegulons that promote glial differentiation (OLIG2, SOX9, NFIA, RFX4). Upon loss of DLX1/DLX2, the quantity and maturity of OPCs increases, whereas differentiation of ventral IPs is stalled, resulting in reduced GABAergic INs. **(C)** DLX2 regulation of Notch signalling in promoting GABAergic IN differentiation primarily takes place in the VZ and SVZ of the GE. In *Dlx1/Dlx2^-/-^* GE, upregulated *Notch1* expression in radial glia in the VZ increases Notch signalling received from the differentiated ventral IPs in the SVZ. Ventral IPs acquire increased *Dll1* and *Dll3* expression, which subsequently increases Notch signalling to neighbouring radial glia and IPs in both the VZ and SVZ. Such increases in Notch signalling therefore arrest IP differentiation and further maturation into GABAergic INs fails.

## Discussion

Our findings show DLX2 transcriptional regulation during neurogenesis controls neuronal-glial cell fate without impacting anatomical lamination of the GE. We uncovered how DLX2 directly inhibits glial fate through direct repression of Notch signalling and glial fate TFs in mid-late neurogenesis, which in turn promotes neuronal differentiation through regional specificity (**Fig. 7B**). Our unbiased spatial approach further established the spatial context of such regulation and expanded our understanding of *Dlx1/Dlx2* regulation of regional specific gene expression in the VZ-SVZ of the GE(Anderson, Qiu, et al., 1997). We identified DLX-independent pathways for GABAergic IN differentiation while also confirming mechanisms of DLX2 function in these processes. These findings collectively define novel mechanisms for DLX2 regulation of neurodevelopmental temporal sequence via a multi-faceted gene regulatory network (**Fig. 7B**).

Studies in the GE and spinal cord have shown that Notch signalling is important in regulating proliferation, differentiation and specification of neural progenitors (Akai et al., 2005; Park & Appel, 2003), and a study published over two decades ago suggested that DLX2 could be upstream of Notch signalling (Yun et al., 2002). Here we demonstrate direct repression of multiple Notch signalling genes by DLX2, including receptors (*Notch1*), ligands (*Dll1*), as well as up- and downstream TFs (*Ascl1, Hes1*), and that such repression is localised to the VZ and SVZ of the developing GE. These findings demonstrate that DLX2 is upstream of the Notch pathway with consequential effects on glial fate repression.

While the upregulation of Notch signalling identified in *Dlx1/Dlx2^-/-^*GE suggests increased cell proliferation, there was only a small increase in number of proliferating progenitors. Instead, the effect of increased Notch signalling was reflected in changes in ventral IPs. *Dlx1/Dlx2^-/-^* ventral IP differentiation was stalled in the early post-mitotic state, coincident with increased Notch signalling, indicating DLX2 repression of Notch signalling is essential for ventral IPs to differentiate into mature GABAergic INs. Cell community and ligand-receptor analysis underscored the importance of cell type and positional regulation by revealing that the effect of upregulated Notch signalling in different cell types was context dependent. Different modes of Notch activation have been shown to impart distinct effects on neural cell differentiation, providing evidence to support the different effects of dysregulated Notch signalling in different regions of the *Dlx1/Dlx2^-/-^* GE(Ables et al., 2010; Basak et al., 2012). Notch signalling through NOTCH1 promotes neural stem cell (NSC) proliferation, thereby maintaining the progenitor pool(Ables et al., 2010). Critically, loss of NOTCH1 affects selectively active NSCs and not quiescent NSCs (Basak et al., 2012). Alternatively, Notch signalling through NOTCH2 and NOTCH3 sustains NSCs quiescence in the adult brain (Engler et al., 2018; Zhang, 2019). Therefore, the enhanced Notch signalling through NOTCH2 and NOTCH3 receptors in *Dlx1/Dlx2^-/-^* ventral progenitors in SVZ communities bordering the VZ (identified as communities 7 and 11) could promote quiescence and slow transition of early progenitors to IPs. Conversely, in the SVZ community that borders the MZ (community 0), Notch signalling through NOTCH1 was enhanced between *Dlx1/Dlx2^-/-^* ventral IPs, promoting proliferation in this cell population, ultimately leading to stalled IP differentiation (**Fig. 7C**). Therefore, repression of Notch signalling by DLX2 in these GE regions (1) allows NSCs to differentiate into IPs, and (2) ensures IP commitment to the GABAergic IN differentiation program.

Despite this stalled differentiation due to upregulated Notch signalling, a proportion of *Dlx1/Dlx2^-/-^* IPs retained the ability to differentiate into mature GABAergic INs and displayed similar transcriptomic profiles to WT GABAergic INs. eRegulon enrichment analysis of *Dlx1/Dlx2^-/-^*ventral IPs suggests that NKX2-1 and/or ISL1 could be key drivers of this DLX1/DLX2 independent pathway, supported by the unaltered *Sst+ Npy+* IN population in *Dlx1/Dlx2^-/-^* GE, which typically arises from *Nkx2-1+* progenitors (Marin et al., 2000). Given the downregulation of other *Sst+ Npy+* INs promoting TFs *Lhx6*, *Lhx8*, and unaltered *Sox6* expression in the *Dlx1/Dlx2^-/-^* GE, this persistent *Sst+ Npy+* IN cell population in the double mutant GE also support that *Lhx6* and *Lhx8* may not be essential for *Sst+ Npy+* INs specification, and activities of *Nkx2-1* and *Sox6* could be sufficient (Asgarian et al., 2022; Munguba et al., 2023). Fundamentally, our findings unravel the complexities of the regulatory networks with DLX1/DLX2-independent processes shown to be partially sufficient for GABAergic IN specification in some IN subtypes. These results also support recent evidence for distinct regulatory functions of combinatorial TF binding to cis-regulatory elements (Catta-Preta et al., 2025), underscoring the complex gene regulatory network that is required for proper telencephalon development.

Increased Notch signalling in *Dlx1/Dlx2^-/-^* impacted neuronal differentiation but also resulted in increased OPC differentiation, aligned with previous studies(Yun et al., 2002). Differentiation trajectory analysis showed increased maturity of *Dlx1/Dlx2^-/-^* OPCs, with increased expression of late OPC markers (*Pdgfra, Sox10*) and lower levels of an early OPC marker (*Cspg4*). Increased activity of SOX5, a regulator of OPC migration, in addition to SOX10, further suggests that *Dlx1/Dlx2^-/-^*OPCs progressed further in the OPC lineage compared to WT OPCs. Additionally, upregulated TCF12 activity in *Dlx1/Dlx2^-/-^* OPCs and ventral NPs also suggest early adoption of OPC fate, consistent with recent mouse model studies showing TCF12 promotes these cell fate decisions(Talley et al., 2023). Downstream of the WNT signalling pathway, TCF7L2 could play a role in mediating postnatal astrocyte maturation in adult mice (Szewczyk et al., 2024), suggesting its potential role in early astroglial specification in *Dlx1/Dlx2^-/-^* GE. The increased OPC signature and differentiation program in distinct cell populations in E12.5-E14.5 *Dlx1/Dlx2^-/-^* GE demonstrates the mechanism by which DLX2 prevents early adoption of OPC fate during neurogenesis.

Our data show that glial differentiation programs driven by NFIA, SOX9, and RFX4 were repressed by DLX2 to inhibit gliogenesis in early progenitors. Upregulated activities of these glial differentiation TFs in *Dlx1/Dlx2^-/-^* progenitors but not in ventral IPs indicate a change in cell fate in ventral progenitors rather than a redirection of ventral IPs into glial cells. These findings establish a novel role for DLX2 in glial fate repression. SOX9 and NFIA are also downstream of Notch signalling and play a role in regulating *Sox2* and *Hes1* expression, respectively(Piper et al., 2010; Wang et al., 2018), further substantiating our finding that DLX2 inhibits gliogenesis via repression of the overall Notch signalling pathway, including its downstream effectors. Intriguingly, NFIA and SOX9 promote both OPC and astrocyte differentiation(Kang et al., 2012; Sim et al., 2009; Stolt et al., 2003). However, astrocytic markers (*Slc1a3, Gfap, Aqp4*) were not upregulated in the *Dlx1/Dlx2^-/-^* GE, suggesting that the increased astroglial TF activity in *Dlx1/Dlx2^-/-^* is not sufficient to promote astrocytic differentiation during mid-neurogenesis (i.e. E12.5-E14.5). Enrichment of astroglial eRegulons in progenitors indicates a degree of commitment to the astroglial lineage in the *Dlx1/Dlx2^-/-^*GE. A potential mechanism may involve the priming of neural progenitors to adopt a glial lineage via glial promoting factors such as SOX9 and NFIA, with subsequent direction towards an oligodendroglial lineage by OPC-promoting TFs, such as OLIG2 and SOX10 (Kang et al., 2012). This illustrates that without OPC differentiating TF activity, glial progenitors will adopt astrocytic fates (Suzuki et al., 2017). Nonetheless, the upregulation of NFIA and SOX9 in *Dlx1/Dlx2^-/-^* GE shown here promoted early glial specification and coupled with elevated OLIG2 and SOX10 activities, accelerated OPC growth, thereby resulting in early oligodendrogenesis.

Spatial transcriptomics at single cell whole genome resolution was essential to characterise the spatial specificity and intricacy of DLX2 regulation of Notch signalling in the ventral IPs and revealed that Notch signalling pathway dysregulation resulted in stalled differentiation in *Dlx1/Dlx2^-/-^*GEs. However, the presence of changes to the OPC differentiation program was not captured by the platform we employed. While a powerful technique for unbiased whole transcriptomic profiling, capture of rare transcripts and therefore of rare cell types such as OPCs presents a technical challenge. Given the specificity required to investigate the small glial populations in the developing telencephalon at E12.5-E14.5, alternative panel-based imaging based spatial profiling methods may enable direct investigation of changes to glial differentiation while still maintaining single-cell resolution.

Our findings cement the role of DLX2 for GABAergic INs specification in developing GE and identified a novel mechanism for inhibiting glial specification during embryonic neurogenesis. DLX2 regulates cell fate decisions in the GE through direct repression of Notch signalling pathway genes in early and intermediate neural progenitors located primarily in the VZ and SVZ. In repressing glial promoting TFs such as SOX9, NFIA, SOX10, and OLIG2, DLX2 inhibits early glial specification, and therefore the growth and maturation of oligodendrocytes at E12.5-E14.5 is blocked. We have therefore identified DLX2 as a glial differentiation repressor via a broad transcriptional regulatory network, redefining the role of DLX2 in forebrain development. In conclusion, we demonstrate a dual role for DLX2 in promoting neurogenesis and repressing gliogenesis, which is necessary for controlling the neurodevelopmental temporal sequence and proper establishment of cell populations in the developing forebrain. Our findings have implications for neurodevelopmental disorders which include paediatric high-grade gliomas, where precursor cells are stalled in differentiation resulting from driver mutations in histone 3 variant genes(Schwartzentruber et al., 2012). The cells-of-origin in diffuse midline glioma (DMG) and diffuse hemispheric gliomas (DHG) have been identified as OPC- and GABAergic IN-like, respectively(Chen et al., 2020; Filbin et al., 2018). Recent studies show that DLX2 knockout reduces tumour burden in DHG-PDX mouse models indicating therapeutic potential in targeting IN cell fate(Liu et al., 2024). Our work now provides mechanistic detail identifying avenues to target neural cell fate that may reduce tumour progression.

## Data Availability

### Lead contact

Further enquiries and requests of information regarding this study should be directed to lead contact David D. Eisenstat.

### Materials availability

All reagents used in the study are available upon request to the lead contact David D. Eisenstat.

### Data and Code availability

Data generated from this study are available with the following GEO accession series: GSE (spatial transcriptomic), GSE292680 (snRNAseq/snATACseq), GSE292442 (bulk ATACseq), GSE292443 (bulk RNAseq), GSE292536 (CUT&RUN & ChIPseq).

## Acknowledgements

This work was supported by an establishment grant from the Royal Children’s Hospital Foundation (2019-1193) to D.D.E. and a BGI STOmics Grant to N.C. and M.R. The Novo Nordisk Foundation Center for Stem Cell Medicine is supported by a Novo Nordisk Foundation grant (NNF21CC0073729). M.R. is funded through a Future Leader Fellowship (107328) from the Heart Foundation of Australia, a Human Frontiers Science Program Grant (RGP008/2024), and a NHMRC Ideas Grant (APP1180905).

## Supplementary Information

Document S1. Figures S1–S7

Table S1. Excel file containing Gene Ontology results from spatial transcriptomics analysis too large to fit in a PDF, related to Figure 1.

Table S2. Excel file containing analysis results of spatial transcriptomics cell community proportions and cell proportion per community too large to fit in a PDF, related to Figure 2.

Table S3. Excel file containing ligand-receptor interactions results from spatial transcriptomics analysis too large to fit in a PDF, related to Figure 2.

Table S4. Excel file containing Gene Ontology results from single nuclei Multiome analysis too large to fit in a PDF, related to Figures 3 and S3.

Table S5. Excel file containing annotation of consensus peaks of DLX2 CUT&RUN, H3K27ac ChIPseq, and ATACseq too large to fit in a PDF, related to Figure 6.

## Author Contributions

Conceptualisation, R.F.L., M.R., M.C.F., and D.D.E.; Methodology, R.F.L., M.C.F., and D.D.E., Software, R.F.L, M.S., and N.C., Visualization, R.F.L., Investigation, R.F.L., A.M.G., P.L.B., Writing – Original Draft, R.F.L., Writing – Review & Editing, R.F.L., M.R., M.C.F., D.D.E., Supervision, M.R., M.C.F., D.D.E., Funding Acquisition, M.R., D.D.E.

## Methods

### Mouse model maintenance

Wildtype (WT) and *Dlx1/Dlx2* double homozygous knockout (*Dlx1/Dlx2^-/-^*) mice(Anderson, Qiu, et al., 1997; Qiu et al., 1995) were maintained in a CD1 background and all animal works conducted were approved by the Murdoch Children’s Research Institute (MCRI) Animal Ethics Committee (AEC). All mouse colonies were housed in the MCRI Disease Modelling Unit in compliance with the MCRI AEC agreements (SPPL20327). Timed pregnant mice were sacrificed by cervical dislocation following MCRI AEC approved protocols at indicated time-points where the presence of a vaginal plug after mating indicated embryonic day 0.5 (E0.5). Genotyping of *Dlx1/Dlx2^-/-^*was performed as previously described (Anderson, Qiu, et al., 1997; Qiu et al., 1995). Embryo dissections were performed in cold PBS, and GE tissue for 10x Multiome were taken from E12.5, E13.5, and E14.5 mice, while GE tissue for ChIPseq, CUT&RUN, RNAseq and ATACseq were taken from E13.5 mice.

### Bulk RNAseq

Total RNA from flash frozen embryonic GE, one *Dlx1/Dlx2^-/-^*and one corresponding WT littermate control from three separate litters, were extracted using the RNeasy Mini kit (Qiagen Cat# 74104). The eluted DNaseq treated RNA was quantified by measuring the absorbance at 260nm, and RNA integrity (RIN) scored determined using the Agilent Tapestation 4150 (Agilent) with RNA Screentapes (Agilent Cat# 5067-5576) and reagents (Agilent Cat# 5067-5577) to ensure a RIN score over 7. The cDNA library construction and sequencing were performed by Australian Genome Research Facility using Novaseq 6000, with 150bp paired reads and targeted recovery of 50 million paired reads per sample.

### Bulk Assay for transposase accessible chromatin with sequencing (ATACseq)

ATACseq was performed using flash frozen embryonic GE, one *Dlx1/Dlx2^-/-^*and one WT control from three separate litters as previously described (Buenrostro et al., 2015). Frozen GEs were homogenised in PBS and lysed in cold lysis buffer (10mM Tris-HCl, pH 7.4 (Thermofisher Scientific Cat# J60636.K2), 10mM NaCl (Sigma Aldrich Cat# S6546), 3mM MgCl_2_ (Thermofisher Scientific Cat# AM9530G), 0.1% (v/v) IGEPAL (Sigma Aldrich Cat# I8896)) for 1 minute, then 50,000 nuclei were taken for the transposition and reaction and purification as previously described (Buenrostro et al., 2015). The purified ATACseq libraries were quantified using the Qubit High Sensitivity dsDNA kit (Thermofisher Scientific Cat# Q32851) and size distribution of the ATACseq libraries was measured using the Tapestation 4150. The sequencing of these libraries was performed at the Victorian Clinical Genomic Services (VCGS) using Novaseq 6000 with 150bp paired-ended reads and targeted recovery of 50 million reads per sample.

### Chromatin Immunoprecipitation with sequencing (ChIPseq) sample preperation

ChIP was performed as previously described (Zhou et al., 2004) using flash frozen WT E13.5 mouse GE. 15 million cells were used for each ChIP reaction, using anti-H3K27ac antibodies (Abcam Cat# ab4729) and anti-IgG antibodies (Thermofisher Scientific Cat# 026102) as a control, and ChIP for each antibody was performed in triplicate. Briefly, tissue was homogenised in cold 1x PBS, then fixed in 1.5% PFA for 20 minutes. The fixed cells were then washed with cold 1x PBS twice, then lysed in ChIP lysis buffer (10mM Tris-HCl, pH 7.4, 10mM EDTA (Thermofisher Scientific Cat# AM9260G), pH 8.0, 1% SDS (Thermofisher Scientific Cat# 15553027), 1x Halt Protease inhibitor (Thermofisher Scientific Cat# 78430)) The chromatin was sheared using an Active Motif EpiShear Sonicator for 15 cycles at 40% pulse strength, with 20 seconds intervals and 50 seconds rest periods. The fragments were then pre-cleared with Pierce Protein A/G Agarose beads (Thermofisher Scientific Cat# 20421) pre-incubated with 0.5 mg/mL BSA for 1 hour at 4°C. The remaining chromatin fragments were separated from beads by centrifugation, then antibodies were added to the chromatin fragments and incubated overnight at 4°C with slight agitation. Pierce Protein A/G Agarose beads were then added to the reaction and incubated for 2 hours at 4°C with slight agitation. Following bead incubation, beads were washed at 4°C with low salt wash buffer (20mM Tris-HCl pH 7.4, 2mM EDTA, pH 8.0, 0.1% SDS, 1% Triton, 150mM NaCl, 1x Halt Protease Inhibitor), high salt wash buffer (20mM Tris-HCl pH 7.4, 2mM EDTA, pH 8.0, 0.1% SDS, 1% Triton, 500mM NaCl, 1x Halt Protease Inhibitor), LiCl buffer (250mM LiCl (Sigma Cat# 62476), 10mM Tris-HCl, pH 7.4, 1mM EDTA, pH 8.0, 1% Sodium Deoxycholate, 1% NP-40, 1x Halt Protease Inhibitor), and twice with 1x TE buffer. Chromatin fragments were then eluted twice with elution buffer (0.1M NaHCO_3_ (Sigma Cat# PHR3591), 1% SDS) pre-heated to 65°C for 15 minutes. Eluted fragments were incubated with 0.01M EDTA, 0.04M Tri-HCl, pH 7.4, 0.1 mg/mL Proteinase K (Promega Cat# MC5008) at 50°C for 2 hours for crosslinking reversal. The eluted DNA fragments were purified using the ChIP DNA Clean and Concentrator (Zymo Research Cat# D5205) according to the manufacturer’s instructions. The purified DNA samples were quantified using the Qubit High Sensitivity dsDNA kit.

### ChIPseq Library Preparation

The ChIPseq library preparation of the H3K27ac and IgG ChIPs were performed using the Illumina TruSeq ChIP Sample Prep Kit (Illumina Cat# IP-202-1012) according to manufacturer’s instructions. The samples were pooled for sequencing at the Walter and Eliza Hall Institute genomics laboratory using Nextseq 2000, with 150 bp single end and targeted recovery of 30 million reads per sample.

### Cleavage Under Target & Release Using Nucleases (CUT&RUN)

CUT&RUN was performed using the CUTANA ChIC/CUT&RUN kit (Epicypher Cat# 14-1048) with slight modifications. Nuclei were extracted using flash frozen WT E13.5 mouse GE from three animals. Frozen tissue was lysed and homogenized in nuclei lysis buffer (10mM Tris-HCl pH 7.4, 10mM NaCl, 3mM MgCl_2_, 0.01% (v/v) Tween-20 (Sigma Aldrich Cat# P1379), 0.01% (v/v) NP-40 (Sigma Aldrich Cat# 74385), 0.001% (w/v) Digitonin (Epicypher Cat# 21-1004), 1% (v/v) BSA (Miltenyi Biotec Cat# 130-091-376), 1x Halt Protease Inhibitor) by trituration, then the lysate was diluted 2-fold with ice cold wash buffer (10mM Tris-HCl pH 7.4, 10mM NaCl, 3mM MgCl_2,_ 1% (v/v) BSA, 0.1% (v/v) Tween-20, 1x Halt Protease Inhibitor). The nuclei were pelleted at 500 rcf for 5 minutes at 4°C, then washed twice and stored in cold wash buffer. 500,00 nuclei were used for each CUT&RUN reaction, using an affinity purified rabbit polyclonal anti-DLX2 antibody(Anderson, Eisenstat, et al., 1997) and an anti-IgG antibody (Epicypher Cat# 13-0042), each performed in triplicates. The final DNA concentration was measured with the Qubit dsDNA HS kit according to the manufacturer’s instructions.

The CUT&RUN libraries were prepared using the CUTANA CUT&RUN Library Prep Kit (Epicypher Cat# 14-1001) according to the manufacturer’s instructions. The concentrations of the final libraries were assessed using Qubit dsDNA HS kit and size fragment distributions were assessed using D1000 Tapestation Reagent and Screentape. Sequencing of the libraries was performed at VCGS using Novaseq X, with 50bp paired-ended read and 5M reads targeted recovery per library.

### Nuclei Extraction, sample processing and library preparation for 10x Multiome

Nuclei were extracted from flash frozen E12.5, E13.5, and E14.5 WT and *Dlx1/Dlx2^-/-^* mouse GE tissues for the 10x Multiome assay. One pair of littermate WT and *Dlx1/Dlx2^-/-^*animals at E12.5 and E14.5 and two pairs of littermate WT and *Dlx1/Dlx2^-/-^* animals at E13.5 were collected. The frozen tissue was homogenized in 0.1X tissue lysis buffer (10mM Tris-HCl pH 7.4, 10mM NaCl, 3mM MgCl_2_, 0.01% (v/v) Tween-20, 0.01% (v/v) NP-40, 0.001% (w/v) Digitonin, 2% (v/v) BSA, 1mM DTT (Sigma Aldrich Cat# 646563), 1U/mL Protector RNase Inhibitor (Sigma Aldrich Cat# 3335399001)) by trituration, then diluted two-folds in cold Single Nuclei Wash Buffer (10mM Tris-HCl pH 7.4, 10mM NaCl, 3mM MgCl_2_, 2% BSA, 0.1% (v/v) Tween-20, 1mM DTT, 1U/mL Protector RNase Inhibitor). The suspension was filtered using a 40μm cell strainer and centrifuged at 500 rcf for 5 minutes at 4°C. The nuclei pellet was washed twice with cold Single Nuclei Wash Buffer, then resuspended in cold diluted Nuclei Buffer (1x Nuclei Buffer (10x Genomics Cat# 2000153), 1mM DTT, 1U/mL Protector RNase Inhibitor).

Following nuclei extraction, samples were processed, and libraries were prepared using 10x Chromium Next GEM Single Cell Multiome ATAC + Gene Expression Reagent kit (10x Genomics Cat # PN-100028) according to 10x Genomics guidelines (CG000338_ChromiumNextGEM_Multiome_ATAC_GEX_User Guide_RevG). E12.5 and E14.5 samples were processed with a targeted recovery of 10,000 nuclei per sample, and E13.5 samples were processed with targeted nuclei recovery of 5,000 nuclei per sample. Nuclei were processed using the Chromium Next GEM Single Cell Multiome Gel Bead Kit A (10x Genomics PN-1000235). Nuclei were loaded into a Chip J (10X Genomics, PN-1000230) and processed on the Chromium Controller (10X Genomics). Subsequent workflow was performed according to the 10x Genomics Chromium Next GEM Single Cell Multiome ATAC + Gene Expression User Guide (CG000338_ChromiumNextGEM_Multiome_ATAC_GEX_User Guide_RevG) at VCGS. The sequencing of the 10x Multiome ATAC and Gene Expression libraries was performed at VCGS using NovaSeq 6000, with a read depth recovery aimed at 412M paired-end reads per sample per library type.

### Stereo-seq Permeabilization Test

The samples were prepared and processed in accordance with manufacturer’s instructions (BGI STOmics Stereo-seq Permeabilization set for Chip-on-a-slide user manual version B) using the STOmics Permeabilisation kit (STOmics Cat# 111KP118). E13.5 mouse heads were fresh frozen in OCT (SAKURA Cat# 4583), cryo-sectioned and mounted onto the Stereoseq P-slide. The ssDNA-stained sections were imaged at 10x magnification to assess permeabilization results, where 15 minutes of permeabilisation was determined as the optimal duration.

### Stereo-seq RNA extraction and cDNA collection

Samples were prepared and processed in accordance with manufacturer’s instructions (BGI STOmics Stereo-seq Transcriptomics set for Chip-on-a-slide user manual version B) using the STOmics Transcriptomics kit (STOmics Cat# 111KT114). Briefly, fresh E12.5, E13.5, and E14.5 WT and *Dlx1/Dlx2^-/-^*mouse heads were embedded and frozen in OCT on an ethanol slurry. Tissues were sectioned at 10μm thick on a CM3050S cryostat (Leica), mounted on a room temperature Stereo-seq T-chip and dried at 37°C for 5 minutes. Once dried, sections were fixed using ice cold methanol for 30 minutes at -30°C. Following fixation, the sections were stained using Tissue Staining Solution (0.5% Qubit ssDNA Reagent, 5X SSC, 5% RI) for 5 mins, and was washed using 0.1X SSC/5% RI. Once dried, glycerol was added, and a coverslip was mounted onto the chip. The stained sections were imaged using the Z2 Axio Imager at 10x magnification.

Following imaging, the sections were permeabilized at 37°C for 15 minutes by incubating with 1 x PR solution (STOmics Cat# 1000028500), followed by a wash step with PR rinse buffer (STOmics Cat# 1000042897) with 5% RI. The sections were then incubated in RT mix (RT reagent (STOmics Cat# 1000042898), RT Oligo (STOmics Cat# 1000028508), RT Additive (STOmics Cat# 1000028502), ReverseT Enzyme (STOmics Cat# 1000042899)) at 42°C for 3 hours for the reverse transcription reaction, followed by incubation in TR buffer (STOmics Cat# 1000028505) at 55°C for 10 mins to remove the tissue section from the chip. To ensure complete removal of the tissue section, the chip was washed twice with 0.1X SSC. Then, cDNA Release Mix (cDNA Release Enzyme (STOmics Cat# 1000028511), cDNA Release Buffer (STOmics Cat# 1000028512)) was added to the chip and incubated at 55°C for 16-18 hrs. cDNA was then collected for purification and library preparation.

The collected cDNA was purified using AMPure XP magnetic beads, with a cDNA:bead ratio of 1:0.8 to select for cDNA fragments larger than 100bp. The purified cDNA was then amplified with the PCR Mix (cDNA Amplification Mix (STOmics Cat# 1000028514), cDNA primer (STOmics Cat# 1000028513)) using the following cycling conditions: 1 cycle at 95°C for 5 minutes, 15 cycles at 98°C for 20 seconds, 58°C for 20 seconds, 72°C 3 minutes, and 1 cycle at 72°C for 5 minutes. The concentration of the amplified cDNA was measured using a Qubit dsDNA HS kit. The amplified cDNA product was further purified using AMPure XP magnetic beads with cDNA:beads ratio of 1:0.6 to remove small (<200bp) cDNA fragments.

To prepare the sequencing library, the pure amplified cDNA products were fragmentated using the Fragmentation Reaction Mix (TMB (STOmics Cat# 1000028517), 0.1x TME (STOmics Cat# 1000028515)) at 55°C for 10 minutes, and the reaction immediately terminated using the Stop Buffer (STOmics Cat# 1000028516). The fragmentated products were amplified using the PCR Amplification Mix (STOmics Cat# 1000029181) with PCR Barcode Primer Mix using the following cycling conditions: 1 cycle at 95°C 5min, 13 cycles at 98 °C 20sec, 58 °C 20sec, 72°C 3mins, and 1 cycle at 72°C 5 mins. The size of the amplified cDNA fragments was measured on the Tapestation 4150 using the HS D5000 Tapestation Reagents (Agilent Cat# 5067-5593) and Screentapes (Agilent Cat# 5067-5592), and the concentration measured with a Qubit dsDNA HS kit. Double sided DNA selection was then performed using AMPure XP magnetic beads, starting with cDNA:beads ratio of 1:0.55 followed by 1:0.7, to select for cDNA sizes of 300-600bp. Quality of the final cDNA library size was assessed using D1000 Tapestation Reagent and Screentape and the concentration measured with Qubit dsDNA HS kit. The sequencing was then performed at the South Australian Genomics Centre using a DNBSEQ T7 sequencer, with 1260 million reads targeted to be recovered per sample.

### Amplification of Dll1 and Ascl1 regulatory regions from gDNA

Regulatory elements of *Notch1* (Chr2:26486674-26486873, Chr2:26501696-26502805), *Dll1* (Chr17:15377751-15378835), and *Ascl1* (Chr10:87494094-87495274) were subcloned into the pGL3 Basic vector that expresses *firefly luc+* (Promega Cat# E1751). For both *Notch1* regulatory regions, the pGL3 Basic vectors containing these regions were synthesized by GenScript (i.e. pGL3_Notch1_1, pGL3_Notch1_2). For *Dll1* and *Ascl1* regulatory regions, primers were designed to amplify these regions from mouse genomic DNA, including *Xho*I and *Kpn*I restriction sites at the 5’ ends of primers, which were used for subcloning into the pGL3 Basic vector.

Genomic DNA was extracted from WT E13.5 embryonic mouse tails and used for PCR containing 1x KAPA HiFi HotStart Ready Mix, 0.3µM of either primer pair for *Dll1* (5’ → 3’ GGTACCTCATCTGTGTTGGGCCATCT, CTCGAGTCCCCGTTAGCTCTGTGTTT) or *Ascl1* (5’ → 3’ GGTACCCAGGGAAGGGTTTAGGCAGA, CTCGAGAGATTGTCAAGAGGCCAGCT), with the following program: 1 cycle at 95°C for 3 minutes, 30 cycles at 98°C for 20 seconds, 59°C for 15 seconds, and 72°C for 1 minute, and 1 cycle at 72°C for 1 minute. The PCR products were then separated on a 1.0% (w/v) agarose gel and the amplified fragments of the predicted size were extracted and purified using the Monarch DNA Gel Extraction Kit (New England Biolabs (NEB) Cat# T1020), according to the manufacturer’s instructions.

The purified DNA products and 5µg of pGL3 Basic vector were digested with 10 units of *Xho*I (NEB Cat# R0146) and 10 units of *Kpn*I-HF (NEB Cat# R3142) enzymes, with 1x rCutSmart buffer (NEB Cat# B6004S). Following incubation at 37°C for 1 hour, the products were separated on a 1% (w/v) agarose gel. The DNA fragments of the corresponding sizes were extracted and purified using the Monarch DNA Gel Extraction kit according to the manufacturer’s guidelines and the concentration of the purified DNA products was determined by measuring the absorbance of the sample at A_260_.

### Ligation of Dll1 and Ascl1 regulatory regions with pGL3 Luciferase reporter vector

To ligate the inserts into the pGL3 Basic vector, 75.55ng of *Dll1* insert and 85.72ng of *Ascl1* insert were used to achieve a 7:1 insert to vector (50 ng) molar ratio. Insert and plasmid backbone were mixed with 1x T4 DNA ligase buffer (NEB Cat# B0202S) and 400 units of T4 DNA ligase (NEB Cat# M0202S), and incubated at room temperature for 10 minutes, followed by heat inactivation at 65°C for 10 minutes. Ligated products (pGL3_Dll1 and pGL3_Ascl1) were transformed into DH5α high competent *E. Coli* (Thermofisher Scientific Cat# EC0112).

### Transformation of pGL3_Dll1 and pGL3_Ascl1 into DH5α E. Coli

DH5α high competent *E. Coli* cells were thawed on ice, and 2µL of the ligation products were added to the cell suspension and incubated on ice for 30 minutes. The cells and plasmid mixture were incubated at 42°C for 30 seconds, then on ice for 2 minutes. Following the addition of LB media after the incubation, the cells were incubated at 37°C for 2 hours with agitation, then plated on LB agar plates containing 1 mg/mL ampicillin (Sigma Aldrich Cat# A1593) and incubated at 37°C for 16-18 hours.

Following incubation, 12 individual colonies were picked and dissolved in 5µL milliQ H_2_O, which was used for colony screen PCR using primers specific to the *Dll1* or *Ascl1* inserts. The remainder of the picked colony was stored in 2mL of LB media containing 1 mg/mL ampicillin. The PCR contained 1x Standard *Taq* Reaction Buffer, 200µM dNTPs, 0.2 µM of each primer, 0.625 units of *Taq* polymerase, and the 5µL of template DNA from the bacterial colonies. The following reaction program was used: 1 cycle at 95°C for 30 seconds, 30 cycles at 95°C for 15 seconds, 59°C for 1 minute, and 68°C for 1 minute, and 1 cycle at 72°C of 5 minutes, and products were then separated on a 1% (w/v) agarose gel. Plasmid DNA of positively screened colony was then prepared using the ZymoPure II Plasmid Midiprep kit (Zymo Research Cat# D4201), according to the manufacturer’s instructions. Final plasmid concentrations were determined by measuring the absorbance of the sample at A_260_.

### Maintenance of human embryonic kidney 293 transformed (HEK293T) cells

HEK293T cells were obtained at MCRI and were cultured in Dulbecco’s Modified Eagle’s Media (DMEM) with 10% FBS (MCRI Tissue Culture) and grown in a 37°C, 5% CO_2_ humidified incubator. Cells were passaged once every four days, where media was discarded, and cells washed with 1xPBS once. Cells were incubated in 0.025% trypsin/EDTA (1mL) (MCRI Tissue Culture) at 37°C, 5% CO_2_ for 5 minutes, then collected in DMEM with 10% FBS (9mL). 1:40 cells were then reseeded into tissue culture flasks and incubated at 37°C with 5% CO_2_.

### Dual Luciferase Reporter Assay with pGL3_Dll1, pGL3_Notch1, and pGL3_Ascl1

pGL3 control vectors and pGL3 vectors containing subcloned promoter regions were transfected into HEK293T cells along with functional mouse DLX2 expressed in pcDNA3 (pcDNA3_DLX2). HEK293T (4×10^3^) cells were seeded per well of a 96-well plate, cultured in DMEM with 10% FBS (MCRI Tissue Culture) and incubated for 24 hours. pGL3 vectors (pGL3_Basic, pGL3_Notch1_1, pGL3_Notch1_2, pGL3_Dll1, pGL3_Ascl1) (10ng), pcDNA3 vectors (pcDNA3, pcDNA3_DLX2) (20ng), pRL (0.5ng) (Promega Cat# E2231) (Table 2.10), and FuGENE transfection reagent at a ratio of 3µL per µg of DNA (Promega Cat# E2311) were added to OPTI-MEM (MCRI Tissue Culture) in a final volume of 5 µL and the mixture incubated at room temperature for 5 minutes. The DNA:FuGENE complex was added to the HEK293T cells and incubated at 37°C, 5% CO_2_ for 72 hours.

Following incubation, the transfected cells were harvested using the Dual-Luciferase Reporter Assay System kit (Promega Cat #E1910), according to the manufacturer’s instructions. Briefly, Passive Lysis Buffer was added to the transfected HEK293T cells and incubated for 15 minutes at room temperature with gentle agitation. The cell lysates were then transferred to a white, flat bottom 96-well plate, and the Luciferase and *Renilla* luminescence were measured by addition of 50µL of 0.2x Luciferase Assay Reagent followed by 50µL 0.2x Stop & Glo Reagent from the Dual-Luciferase Reporter Assay System kit (Promega Cat# E1910), respectively. Data analysis, visualisation and statistical analysis were performed using Prism Graph Pad (v.10), where a non-parametric Mann Whitney test was used to calculate statistical significance.

## Bioinformatics analysis of multiomic sequencing based data

### Pre-processing and differential expression analysis of bulk RNAseq data

The raw RNAseq data from WT and *Dlx^-^/Dlx2^-/-^* samples were pre-processed using the RNAsik pipeline (v1.5.5) (Tsyganov et al., 2018) to produce a raw genes count matrix and calculate quality control metrics. Raw counts were used for downstream differential expression analysis using the Degust web tool (Powell, 2019), which subsequently produced plots outlining quality metrics including multidimensional scaling and MA plots. The differential expression analysis was computed using Limma-voom workflow with counts per million library size normalization. By thresholding the false discovery rate (FDR) at <0.05, a list of differentially expressed genes (DEGs) was obtained, and gene ontology and pathway analyses were subsequently performed using the Metascape webtool (Zhou et al., 2019) with default settings.

### Pre-processing and analysis of bulk ATACseq data

Raw sequencing reads for all ATACseq samples were processed through the nf-core/atacseq pipeline (v1.2.1) (Ewels et al., 2020; Patel et al., 2020). The GRCm38/mm10 genome was used for the sequence alignment performed using BWA; and the peaks called using MACS2 were then quantified using featureCounts to create a raw peak count matrix and used for differential chromatin accessibility analysis using the Degust webtool, with a threshold of significance at FDR<0.05. The peaks that were differentially accessible were then annotated using the Genomic Regions Enrichment of Annotations Tool (GREAT) (McLean et al., 2010), with the basal plus extension setting of 5kb proximal, 1kb downstream and up to 1000kb distal. Custom R scripts were used to perform gene set enrichment analysis, using clusterProfiler (v4.2.2) (Wu et al., 2021) and ReactomePA (v1.38.0) (Yu & He, 2016) with standard configurations, and results plotted with ggplot2 (v3.5.0). The visualization of peaks was then performed using the University of California Santa Cruz (UCSC) genome browser (Perez et al., 2024).

### Pre-processing and analysis of ChIPseq and CUT&RUN data

The raw sequencing results of the ChIPseq and CUT&RUN were processed through the nf-core/chipseq (v1.2.2) (Patel et al., 2021) and nf-core/cutandrun (v3.2.2) (Cheshire et al., 2024) pipeline, respectively, to generate peaks sets and assess the quality of the raw sequencing results. BWA was used for the sequence alignment of the ChIPseq pipeline and Bowtie2(Langmead & Salzberg, 2012) was used for the CUT&RUN pipeline, with both using the GRCm38/mm10 mouse genome from ENSEMBL as a reference. Bigwig files were generated for visualisation and peaks were called against the IgG samples using MACS2. The consensus peak sets of each replicate, and the consensus of H3K27ac, ATACseq, and DLX2 CUT&RUN were performed using Bedtools intersect (Quinlan & Hall, 2010). The genomic region annotation of these peak sets was then performed in R (v4.1.2) using clusterProfiler (v4.2.2), while the gene association and subsequent gene set enrichment analysis was performed using GREAT. The set association was performed with the basal plus extension setting of 5kb proximal, 1kb downstream and up to 1000kb distal. HOMER (v4.1.1) (Heinz et al., 2010) was then used to perform motif enrichment analysis, using findMotifsGenome.pl with GRCm38/mm10 genome and ‘size’ parameter set as 100.

### Pre-processing and downstream analyses of 10x Multiome data

The raw sequencing fastq files of the E12.5, E13.5, and E14.5 WT and *Dlx1/Dlx2^-/-^* Gene Expression and ATAC libraries were processed using Cell Ranger Arc (v2.0.2), specifically the cellranger-arc count function with default parameters. The mm10 reference genome was taken from 10x Genomics and was used for all analyses.

### Pre-processing analysis of snRNAseq data

Downstream gene expression data pre-processing was performed in Scanpy (v1.10.1)(Wolf et al., 2018) with individual output gene expression count matrix (filtered_feature_bc_matrix.h5) from the cellranger-arc count pipeline as input. The data was first filtered for doublet detection using scanpy.pp.scrublet, then for cells with less than 8000 genes, less than 30,000 counts, and less than 5% mitochondrial gene reads. The individual data sets were then concatenated to create a merged data set. Read counts of the merged data were normalized and log transformed, and the highly variable genes were calculated with min_mean=0.0125 and max_mean=3, and min_disp=0.5. The data was then filtered for these highly variable genes, followed by principal component analysis (PCA) using the ARPACK SVD solver. The individual data sets were integrated using scanpy.external.pp.harmony_integrate, and the neighbours were calculated using the integrated PCA with n_neighbours=10 and n_pcs=50. The uniform manifold approximation and projection (UMAP) was calculated using default parameters, and Leiden unsupervised clustering was performed. The clusters were then manually annotated by examining differentially overexpressed genes in each cluster, as well as using a list of marker genes for various expected cell types in the GE at E12.5 to E14.5, including radial glia (*Vim, Sox2. Mki67, Top2a*), ventral neural progenitors (*Ascl1, Gli3, Adgrv1*), ventral intermediate progenitors (*Dcx, Nkx2-1, St18*), oligodendroglial precursor cells (*Cspg4, Olig2, Sox10*, *Pdgfra*), GABAergic interneurons (*Gad1, Gad2, Nrxn3*, *Slc32a1*), *Stomatostatin/Neuropeptide-Y* expressing interneurons (*Sst, Npy, Sox6, Erbb4)*, *Gad* negative interneurons (*Nrxn1, Ank3)*, dorsal intermediate progenitors (*Tbr1, Eomes*), and glutamatergic neurons (*Slc1a7, Grin2b*).

### Pre-processing analysis of snATACseq data

Downstream ATAC pre-processing was performed using pycisTopic (v1.0.3), where the WT and the *Dlx1/Dlx2^-/-^*data were processed separately, and the cellranger-arc count ATAC fragment outputs (atac_fragments.tsv.gz) were individually loaded into pycisTopic. A range of cell quality filtering was performed, including determining the total number of unique fragments, enrichment at transcription start sites (TSS), and fraction of reads in peaks (FRIP). The filtering threshold was then computed using pycisTopc.qc with automatic detection and the data set filtered with corresponding parameters. Using the annotated snRNAseq data generated previously as a reference, pseudobulk ATAC fragment profiles for each cell type were then generated and peaks were called using MACS for each pseudobulk profile, followed by inferring a consensus peak set for each cell type. cisTopic object was then created for the individual data sets, including filtering of GRCm38/mm10 genome blacklist regions, obtained from the UCSC database. cisTopic objects were merged according to genotype, i.e. the WT data were merged into one object and *Dlx1/Dlx2^-/-^* data into another. The WT and *Dlx1/Dlx2^-/-^* data sets were individually integrated using Harmony. Candidate enhancer regions were then inferred by binarization of region-topic probability, and differential accessible regions of each cell type was computed within the same genotype.

### Differentiation trajectory analysis of snRNAseq data

The snRNAseq differentiation trajectory of the ventrally derived cell types, including radial glia, ventral NPs, ventral IPs, OPCs, GABAergic INs, and *Sst+ Npy+* INs, were computed using scFates (v1.0.7) (Faure et al., 2023). The scRNAseq was first filtered for targeted cell types, and multiscale diffusion space was computed using Palantir (v1.3.3)(Setty et al., 2019), and neighbours were subsequently computed in this space. ForceAtlas2 embedding was then calculated using the first 2 principal components as initial positions, and the differentiation tree was determined using scFates.tl.tree with the simpleppt option, using 200 nodes with the following parameters: lambda 180, sigma 0.075, and nsteps 200. The pseudotime and dendrogram of the differentiation tree were then calculated using scFates.tl.pseudotime and scFates.tl.dendrogram, respectively.

### Enhancer driven gene regulatory network analysis

The SCENIC+ (v1.01a1)(Bravo Gonzalez-Blas et al., 2023) pipeline was used to identify enhancer driven gene regulatory networks (eRegulons) in WT and *Dlx1/Dlx2^-/-^* snMultiome data. The pre-processed snRNAseq data was first separated into an anndata object containing either WT or *Dlx1/Dlx2^-/-^*data, which was then used for the analysis of the data sets in each genotype, respectively. The pre-processing ATAC data as the cisTopic object was used as snATACseq data input, along with the topic set that was generated from the pycisTopic processing.

pycisTarget (v1.0.2) (Bravo Gonzalez-Blas et al., 2023) was used to generate custom databases of scores of a set of given motifs in the regulatory regions identified in the data set. Separate data bases for the WT and *Dlx1/Dlx2^-/-^*data were generated using create_cistarget_motif_databases, where the consensus peak regions previously generated from pycisTopic were used as input. The list of mm10 motifs and transcription factors expressed in mice were obtained from the cisTarget database, which was also used for the SCENIC+ pipeline (https://resources.aertslab.org/cistarget/). The eRegulon specificity score was determined using scenicplus.RSS.regulon_specificity_scores function, with variable set as cell types to compute the eRegulon specificity score per cell type. The result table was then exported and plotted in Python using matplotlib (v3.6.3)(Hunter, 2007).

### snRNAseq Pseudobulk differential expression and transcription factor activity analysis

The pseudobulk profile of the snRNAseq data was obtained using Decoupler (v1.6.0)(Badia et al., 2022) get_pseudobulk function, with rounded raw counts and counts grouped by cell types. The read counts of each cell type were extracted, and the genes were filtered using decoupler.filter_by_expr, with min_count=10 and min_total_count=15. The WT vs *Dlx1/Dlx2^-/-^* differential expression of the filtered genes were then computed using pydeseq2 (v0.4.4)(Muzellec et al., 2023). The differential gene expression matrix of each cell type was then extracted for downstream usage.

To examine the transcription factor activity changes, the differential expression fold changes were collated with the SCENIC+ results. The WT and *Dlx1/Dlx2^-/-^* SCENIC+ results generated a matrix containing a list of gene targets for each TF, and the corresponding transcriptional effect, i.e. either positive regulation or negative regulation. These were extracted from the analysis of each genotype and combined to create one TF-gene target network. Along with the differential gene expression matrix from each cell type these were used as input for the decoupler.run_ulm function to compute the TF activity scores in each cell type. The networks were visualized using decoupler.plot_network.

### Pre-processing analysis of spatial transcriptomics data

The stitched images taken during tissue processing were processed through ImageStudio to transform into files compatible with the SAW pipeline. The paired fastq, mask and processed imaging files (.ipr and.tar.gz) were then processed using the SAW pipeline (v6.12) using default parameters including cell binning(Gong et al., 2024). Files generated from mouse heads were aligned to the reference genome GRCm39.

Downstream analysis was performed in Stereopy (v1.4.0)(Fang et al., 2023). Respective tissue.gef output files from the SAW pipeline were loaded into Stereopy. The E12.5, E13.5, and E14.5 mouse head data were analysed at bin size 20, and were filtered to have at least 100 counts, 50 genes and no more than 5% of mitochondrial genes. Genes were only included if expressed in at least 15 cells. The data was normalised and log transformed, followed by PCA using 50 principal components and ARPACK SVD solver. Neighbours were calculated followed by gaussian smoothing calculated with n_neighbours=10. The data was then exported into Scanpy and Squidpy compatible anndata objects.

### Spatial transcriptomics analysis

Individually, the pre-processed data were loaded into Squidpy (v1.2.3)(Palla et al., 2022), where neighbours were calculated using n_neighbors=10 and n_pcs=20, and UMAP and Leiden clustering were subsequently performed. Clusters that corresponded to the mice forebrain were extracted. The data was next loaded into Stereopy(Fang et al., 2023) creating a multi-slice object. With this combined data set, PCA was performed using 50 principal components and then the data integrated using stereopy.tl.batches_integrate. Then neighbours and spatial neighbours were computed, and UMAPs were calculated using both neighbours and spatial neighbours. Leiden clustering of each UMAP result was performed, and marker genes identified using stereopy.tl.find_marker_genes. Using the identified marker genes, tissue locations, and a list of selected anatomical region markers, the clusters were manually annotated. These included ventricular zone/neural progenitors (*Fabp7, Nes*), subventricular zone/intermediate progenitors (*Ascl1, St18*), mantle zone/neurons (*Dcx, Tubb3*), lateral ganglionic eminence (*Gsx2, Pak3*), medial ganglionic eminence (*Nkx2-1, Olig2*), and neocortex (*Tbr1, Eomes, Pax6*). Corresponding dot plot figures were generated using stereopy.pl.cluster_gene_scatter.

The cell communities were computed using stereopy.tl.ms_community_detection and the agglomerative algorithm, with the following parameters: annotation=cell types, window size=150, sliding steps=50, resolution=0.2, scatter threshold=0.12, and number of clusters=16.

To perform cell communication analysis within the detected cell communities, the individual cell community anndata of each timepoint and genotype were imported into Stereopy. The cell communication analysis was then performed using stereopy.tl.cell_cell_communication with the Liana database(Dimitrov et al., 2022). The cell communication analysis results of all timepoints in WT and *Dlx1/Dlx2^-/-^* were combined for downstream visualisation.

## References

Ables, J. L., Decarolis, N. A., Johnson, M. A., Rivera, P. D., Gao, Z., Cooper, D. C., Radtke, F., Hsieh, J., & Eisch, A. J. (2010). Notch1 is required for maintenance of the reservoir of adult hippocampal stem cells. J Neurosci, 30(31), 10484–10492. 10.1523/JNEUROSCI.4721-09.2010

Akai, J., Halley, P. A., & Storey, K. G. (2005). FGF-dependent Notch signaling maintains the spinal cord stem zone. Genes Dev, 19(23), 2877–2887. 10.1101/gad.357705

Anderson, S. A., Eisenstat, D. D., Shi, L., & Rubenstein, J. L. (1997). Interneuron migration from basal forebrain to neocortex: dependence on Dlx genes. Science, 278(5337), 474–476. 10.1126/science.278.5337.474

Anderson, S. A., Kaznowski, C. E., Horn, C., Rubenstein, J. L., & McConnell, S. K. (2002). Distinct origins of neocortical projection neurons and interneurons in vivo. Cereb Cortex, 12(7), 702–709. 10.1093/cercor/12.7.702

Anderson, S. A., Qiu, M., Bulfone, A., Eisenstat, D. D., Meneses, J., Pedersen, R., & Rubenstein, J. L. (1997). Mutations of the homeobox genes Dlx-1 and Dlx-2 disrupt the striatal subventricular zone and differentiation of late born striatal neurons. Neuron, 19(1), 27–37. 10.1016/s0896-6273(00)80345-1

Asgarian, Z., Oliveira, M. G., Stryjewska, A., Maragkos, I., Rubin, A. N., Magno, L., Pachnis, V., Ghorbani, M., Hiebert, S. W., Denaxa, M., & Kessaris, N. (2022). MTG8 interacts with LHX6 to specify cortical interneuron subtype identity. Nat Commun, 13(1), 5217. 10.1038/s41467-022-32898-6

Badia, I. M. P., Velez Santiago, J., Braunger, J., Geiss, C., Dimitrov, D., Muller-Dott, S., Taus, P., Dugourd, A., Holland, C. H., Ramirez Flores, R. O., & Saez-Rodriguez, J. (2022). decoupleR: ensemble of computational methods to infer biological activities from omics data. Bioinform Adv, 2(1), vbac016. 10.1093/bioadv/vbac016

Bandler, R. C., Mayer, C., & Fishell, G. (2017). Cortical interneuron specification: the juncture of genes, time and geometry. Curr Opin Neurobiol, 42, 17–24. 10.1016/j.conb.2016.10.003

Baroti, T., Zimmermann, Y., Schillinger, A., Liu, L., Lommes, P., Wegner, M., & Stolt, C. C. (2016). Transcription factors Sox5 and Sox6 exert direct and indirect influences on oligodendroglial migration in spinal cord and forebrain. Glia, 64(1), 122–138. 10.1002/glia.22919

Basak, O., Giachino, C., Fiorini, E., Macdonald, H. R., & Taylor, V. (2012). Neurogenic subventricular zone stem/progenitor cells are Notch1-dependent in their active but not quiescent state. J Neurosci, 32(16), 5654–5666. 10.1523/JNEUROSCI.0455-12.2012

Bravo Gonzalez-Blas, C., De Winter, S., Hulselmans, G., Hecker, N., Matetovici, I., Christiaens, V., Poovathingal, S., Wouters, J., Aibar, S., & Aerts, S. (2023). SCENIC+: single-cell multiomic inference of enhancers and gene regulatory networks. Nat Methods, 20(9), 1355–1367. 10.1038/s41592-023-01938-4

Buenrostro, J. D., Wu, B., Chang, H. Y., & Greenleaf, W. J. (2015). ATAC-seq: A Method for Assaying Chromatin Accessibility Genome-Wide. Curr Protoc Mol Biol, 109, 21 29 21–21 29 29. 10.1002/0471142727.mb2129s109

Cadwell, C. R., Bhaduri, A., Mostajo-Radji, M. A., Keefe, M. G., & Nowakowski, T. J. (2019). Development and Arealization of the Cerebral Cortex. Neuron, 103(6), 980–1004. 10.1016/j.neuron.2019.07.009

Catta-Preta, R., Lindtner, S., Ypsilanti, A., Seban, N., Price, J. D., Abnousi, A., Su-Feher, L., Wang, Y., Cichewicz, K., Boerma, S. A., Juric, I., Jones, I. R., Akiyama, J. A., Hu, M., Shen, Y., Visel, A., Pennacchio, L. A., Dickel, D. E., Rubenstein, J. L. R., & Nord, A. S. (2025). Combinatorial transcription factor binding encodes cis-regulatory wiring of mouse forebrain GABAergic neurogenesis. Dev Cell, 60(2), 288–304 e286. 10.1016/j.devcel.2024.10.004

Chen, A., Liao, S., Cheng, M., Ma, K., Wu, L., Lai, Y., Qiu, X., Yang, J., Xu, J., Hao, S., Wang, X., Lu, H., Chen, X., Liu, X., Huang, X., Li, Z., Hong, Y., Jiang, Y., Peng, J.,…Wang, J. (2022). Spatiotemporal transcriptomic atlas of mouse organogenesis using DNA nanoball-patterned arrays. Cell, 185(10), 1777–1792 e1721. 10.1016/j.cell.2022.04.003

Chen, C. C. L., Deshmukh, S., Jessa, S., Hadjadj, D., Lisi, V., Andrade, A. F., Faury, D., Jawhar, W., Dali, R., Suzuki, H., Pathania, M., A, D., Dubois, F., Woodward, E., Hebert, S., Coutelier, M., Karamchandani, J., Albrecht, S., Brandner, S.,…Jabado, N. (2020). Histone H3.3G34-Mutant Interneuron Progenitors Co-opt PDGFRA for Gliomagenesis. Cell, 183(6), 1617–1633 e1622. 10.1016/j.cell.2020.11.012

Cheshire, C., Charlotte-west, Rönkkö, T., bot, n.-c., Patel, H., tamara-hodgetts, Ladd, D., Thiery, A., Fields, C., Deu-Pons, J., Ewels, P., Möller, S., & Menden, K. (2024). nf-core/cutandrun: nf-core/cutandrun v3.2.2 Iridium Ibis. In Zenodo.

Chillakuri, C. R., Sheppard, D., Lea, S. M., & Handford, P. A. (2012). Notch receptor-ligand binding and activation: insights from molecular studies. Semin Cell Dev Biol, 23(4), 421–428. 10.1016/j.semcdb.2012.01.009

Cobos, I., Borello, U., & Rubenstein, J. L. (2007). Dlx transcription factors promote migration through repression of axon and dendrite growth. Neuron, 54(6), 873–888. 10.1016/j.neuron.2007.05.024

Dimitrov, D., Turei, D., Garrido-Rodriguez, M., Burmedi, P. L., Nagai, J. S., Boys, C., Ramirez Flores, R. O., Kim, H., Szalai, B., Costa, I. G., Valdeolivas, A., Dugourd, A., & Saez-Rodriguez, J. (2022). Comparison of methods and resources for cell-cell communication inference from single-cell RNA-Seq data. Nat Commun, 13(1), 3224. 10.1038/s41467-022-30755-0

Engler, A., Zhang, R., & Taylor, V. (2018). Notch and Neurogenesis. Adv Exp Med Biol, 1066, 223–234. 10.1007/978-3-319-89512-3_11

Ewels, P. A., Peltzer, A., Fillinger, S., Patel, H., Alneberg, J., Wilm, A., Garcia, M. U., Di Tommaso, P., & Nahnsen, S. (2020). The nf-core framework for community-curated bioinformatics pipelines. Nat Biotechnol, 38(3), 276–278. 10.1038/s41587-020-0439-x

Fang, S., Xu, M., Cao, L., Liu, X., Bezulj, M., Tan, L., Yuan, Z., Li, Y., Xia, T., Guo, L., Kovacevic, V., Hui, J., Guo, L., Liu, C., Cheng, M., Lin, L. a., Wen, Z., Josic, B., Milicevic, N.,…Xu, X. (2023). Stereopy: modeling comparative and spatiotemporal cellular heterogeneity via multi-sample spatial transcriptomics. bioRxiv, 2023.2012.2004.569485. 10.1101/2023.12.04.569485

Farah, E. N., Hu, R. K., Kern, C., Zhang, Q., Lu, T. Y., Ma, Q., Tran, S., Zhang, B., Carlin, D., Monell, A., Blair, A. P., Wang, Z., Eschbach, J., Li, B., Destici, E., Ren, B., Evans, S. M., Chen, S., Zhu, Q., & Chi, N. C. (2024). Spatially organized cellular communities form the developing human heart. Nature, 627(8005), 854–864. 10.1038/s41586-024-07171-z

Faure, L., Soldatov, R., Kharchenko, P. V., & Adameyko, I. (2023). scFates: a scalable python package for advanced pseudotime and bifurcation analysis from single-cell data. Bioinformatics, 39(1). 10.1093/bioinformatics/btac746

Filbin, M. G., Tirosh, I., Hovestadt, V., Shaw, M. L., Escalante, L. E., Mathewson, N. D., Neftel, C., Frank, N., Pelton, K., Hebert, C. M., Haberler, C., Yizhak, K., Gojo, J., Egervari, K., Mount, C., van Galen, P., Bonal, D. M., Nguyen, Q. D., Beck, A.,…Suva, M. L. (2018). Developmental and oncogenic programs in H3K27M gliomas dissected by single-cell RNA-seq. Science, 360(6386), 331–335. 10.1126/science.aao4750

Fong, B. C., Chakroun, I., Iqbal, M. A., Paul, S., Bastasic, J., O’Neil, D., Yakubovich, E., Bejjani, A. T., Ahmadi, N., Carter, A., Clark, A., Leone, G., Park, D. S., Ghanem, N., Vandenbosch, R., & Slack, R. S. (2022). The Rb/E2F axis is a key regulator of the molecular signatures instructing the quiescent and activated adult neural stem cell state. Cell Rep, 41(5), 111578. 10.1016/j.celrep.2022.111578

Gong, C., Li, S., Wang, L., Zhao, F., Fang, S., Yuan, D., Zhao, Z., He, Q., Li, M., Liu, W., Li, Z., Xie, H., Liao, S., Chen, A., Zhang, Y., Li, Y., & Xu, X. (2024). SAW: an efficient and accurate data analysis workflow for Stereo-seq spatial transcriptomics. GigaByte, 2024, gigabyte111. 10.46471/gigabyte.111

Grandbarbe, L., Bouissac, J., Rand, M., Hrabe de Angelis, M., Artavanis-Tsakonas, S., & Mohier, E. (2003). Delta-Notch signaling controls the generation of neurons/glia from neural stem cells in a stepwise process. Development, 130(7), 1391–1402. 10.1242/dev.00374

Heinz, S., Benner, C., Spann, N., Bertolino, E., Lin, Y. C., Laslo, P., Cheng, J. X., Murre, C., Singh, H., & Glass, C. K. (2010). Simple combinations of lineage-determining transcription factors prime cis-regulatory elements required for macrophage and B cell identities. Mol Cell, 38(4), 576–589. 10.1016/j.molcel.2010.05.004

Hunter, J. D. (2007). Matplotlib: A 2D Graphics Environment. Computing in Science & Engineering, 9(3), 90–95. 10.1109/MCSE.2007.55

Jiang, X., & Nardelli, J. (2016). Cellular and molecular introduction to brain development. Neurobiol Dis, 92(Pt A), 3–17. 10.1016/j.nbd.2015.07.007

Kang, P., Lee, H. K., Glasgow, S. M., Finley, M., Donti, T., Gaber, Z. B., Graham, B. H., Foster, A. E., Novitch, B. G., Gronostajski, R. M., & Deneen, B. (2012). Sox9 and NFIA coordinate a transcriptional regulatory cascade during the initiation of gliogenesis. Neuron, 74(1), 79–94. 10.1016/j.neuron.2012.01.024

Langmead, B., & Salzberg, S. L. (2012). Fast gapped-read alignment with Bowtie 2. Nat Methods, 9(4), 357–359. 10.1038/nmeth.1923

Le, T. N., Du, G., Fonseca, M., Zhou, Q. P., Wigle, J. T., & Eisenstat, D. D. (2007). Dlx homeobox genes promote cortical interneuron migration from the basal forebrain by direct repression of the semaphorin receptor neuropilin-2. J Biol Chem, 282(26), 19071–19081. 10.1074/jbc.M607486200

Le, T. N., Zhou, Q. P., Cobos, I., Zhang, S., Zagozewski, J., Japoni, S., Vriend, J., Parkinson, T., Du, G., Rubenstein, J. L., & Eisenstat, D. D. (2017). GABAergic Interneuron Differentiation in the Basal Forebrain Is Mediated through Direct Regulation of Glutamic Acid Decarboxylase Isoforms by Dlx Homeobox Transcription Factors. J Neurosci, 37(36), 8816–8829. 10.1523/JNEUROSCI.2125-16.2017

Lindtner, S., Catta-Preta, R., Tian, H., Su-Feher, L., Price, J. D., Dickel, D. E., Greiner, V., Silberberg, S. N., McKinsey, G. L., McManus, M. T., Pennacchio, L. A., Visel, A., Nord, A. S., & Rubenstein, J. L. R. (2019). Genomic Resolution of DLX-Orchestrated Transcriptional Circuits Driving Development of Forebrain GABAergic Neurons. Cell Rep, 28(8), 2048–2063 e2048. 10.1016/j.celrep.2019.07.022

Liu, I., Alencastro Veiga Cruzeiro, G., Bjerke, L., Rogers, R. F., Grabovska, Y., Beck, A., Mackay, A., Barron, T., Hack, O. A., Quezada, M. A., Molinari, V., Shaw, M. L., Perez-Somarriba, M., Temelso, S., Raynaud, F., Ruddle, R., Panditharatna, E., Englinger, B., Mire, H. M.,…Filbin, M. G. (2024). GABAergic neuronal lineage development determines clinically actionable targets in diffuse hemispheric glioma, H3G34-mutant. Cancer Cell. 10.1016/j.ccell.2024.08.006

Marin, O., Anderson, S. A., & Rubenstein, J. L. (2000). Origin and molecular specification of striatal interneurons. J Neurosci, 20(16), 6063–6076. 10.1523/JNEUROSCI.20-16-06063.2000

McLean, C. Y., Bristor, D., Hiller, M., Clarke, S. L., Schaar, B. T., Lowe, C. B., Wenger, A. M., & Bejerano, G. (2010). GREAT improves functional interpretation of cis-regulatory regions. Nat Biotechnol, 28(5), 495–501. 10.1038/nbt.1630

Mu, L., Berti, L., Masserdotti, G., Covic, M., Michaelidis, T. M., Doberauer, K., Merz, K., Rehfeld, F., Haslinger, A., Wegner, M., Sock, E., Lefebvre, V., Couillard-Despres, S., Aigner, L., Berninger, B., & Lie, D. C. (2012). SoxC transcription factors are required for neuronal differentiation in adult hippocampal neurogenesis. J Neurosci, 32(9), 3067–3080. 10.1523/JNEUROSCI.4679-11.2012

Munguba, H., Nikouei, K., Hochgerner, H., Oberst, P., Kouznetsova, A., Ryge, J., Munoz-Manchado, A. B., Close, J., Batista-Brito, R., Linnarsson, S., & Hjerling-Leffler, J. (2023). Transcriptional maintenance of cortical somatostatin interneuron subtype identity during migration. Neuron, 111(22), 3590-3603 e3595. 10.1016/j.neuron.2023.07.018

Muzellec, B., Telenczuk, M., Cabeli, V., & Andreux, M. (2023). PyDESeq2: a python package for bulk RNA-seq differential expression analysis. Bioinformatics, 39(9). 10.1093/bioinformatics/btad547

Nord, A. S., & West, A. E. (2020). Neurobiological functions of transcriptional enhancers. Nat Neurosci, 23(1), 5–14. 10.1038/s41593-019-0538-5

Palla, G., Spitzer, H., Klein, M., Fischer, D., Schaar, A. C., Kuemmerle, L. B., Rybakov, S., Ibarra, I. L., Holmberg, O., Virshup, I., Lotfollahi, M., Richter, S., & Theis, F. J. (2022). Squidpy: a scalable framework for spatial omics analysis. Nat Methods, 19(2), 171–178. 10.1038/s41592-021-01358-2

Park, H. C., & Appel, B. (2003). Delta-Notch signaling regulates oligodendrocyte specification. Development, 130(16), 3747–3755. 10.1242/dev.00576

Patel, H., Ewels, P., Peltzer, A., Behrens, D., Gabernet, G., Jin, M., mashehu, & Garcia, M. (2020). nf-core/atacseq: nf-core/atacseq v1.2.1 - Iron Centipede. In 10.5281/zenodo.3965985

Patel, H., Wang, C., Ewels, P., Silva, T. C., Peltzer, A., Behrens, D., Garcia, M., Mashehu, Rotholandus, Haglund, S., & Kretzschmar, W. (2021). nf-core/chipseq: nf-core/chipseq v1.2.2 - Rusty Mole. In Zenodo. 10.5281/zenodo.4711243

Perez, G., Barber, G. P., Benet-Pages, A., Casper, J., Clawson, H., Diekhans, M., Fischer, C., Gonzalez, J. N., Hinrichs, A. S., Lee, C. M., Nassar, L. R., Raney, B. J., Speir, M. L., van Baren, M. J., Vaske, C. J., Haussler, D., Kent, W. J., & Haeussler, M. (2024). The UCSC Genome Browser database: 2025 update. Nucleic Acids Res. 10.1093/nar/gkae974

Petryniak, M. A., Potter, G. B., Rowitch, D. H., & Rubenstein, J. L. (2007). Dlx1 and Dlx2 control neuronal versus oligodendroglial cell fate acquisition in the developing forebrain. Neuron, 55(3), 417–433. 10.1016/j.neuron.2007.06.036

Piper, M., Barry, G., Hawkins, J., Mason, S., Lindwall, C., Little, E., Sarkar, A., Smith, A. G., Moldrich, R. X., Boyle, G. M., Tole, S., Gronostajski, R. M., Bailey, T. L., & Richards, L. J. (2010). NFIA controls telencephalic progenitor cell differentiation through repression of the Notch effector Hes1. J Neurosci, 30(27), 9127–9139. 10.1523/JNEUROSCI.6167-09.2010

Powell, D. (2019). drpowell/degust 4.1.1. In Zenodo.

Qiu, M., Bulfone, A., Martinez, S., Meneses, J. J., Shimamura, K., Pedersen, R. A., & Rubenstein, J. L. (1995). Null mutation of Dlx-2 results in abnormal morphogenesis of proximal first and second branchial arch derivatives and abnormal differentiation in the forebrain. Genes Dev, 9(20), 2523–2538. 10.1101/gad.9.20.2523

Quinlan, A. R., & Hall, I. M. (2010). BEDTools: a flexible suite of utilities for comparing genomic features. Bioinformatics, 26(6), 841–842. 10.1093/bioinformatics/btq033

Schwartzentruber, J., Korshunov, A., Liu, X. Y., Jones, D. T., Pfaff, E., Jacob, K., Sturm, D., Fontebasso, A. M., Quang, D. A., Tonjes, M., Hovestadt, V., Albrecht, S., Kool, M., Nantel, A., Konermann, C., Lindroth, A., Jager, N., Rausch, T., Ryzhova, M.,…Jabado, N. (2012). Driver mutations in histone H3.3 and chromatin remodelling genes in paediatric glioblastoma. Nature, 482(7384), 226–231. 10.1038/nature10833

Setty, M., Kiseliovas, V., Levine, J., Gayoso, A., Mazutis, L., & Pe’er, D. (2019). Characterization of cell fate probabilities in single-cell data with Palantir. Nat Biotechnol, 37(4), 451–460. 10.1038/s41587-019-0068-4

Shimojo, H., Ohtsuka, T., & Kageyama, R. (2008). Oscillations in notch signaling regulate maintenance of neural progenitors. Neuron, 58(1), 52–64. 10.1016/j.neuron.2008.02.014

Sim, F. J., Windrem, M. S., & Goldman, S. A. (2009). Fate determination of adult human glial progenitor cells. Neuron Glia Biol, 5(3-4), 45–55. 10.1017/S1740925X09990317

Stolt, C. C., Lommes, P., Sock, E., Chaboissier, M. C., Schedl, A., & Wegner, M. (2003). The Sox9 transcription factor determines glial fate choice in the developing spinal cord. Genes Dev, 17(13), 1677–1689. 10.1101/gad.259003

Suzuki, N., Sekimoto, K., Hayashi, C., Mabuchi, Y., Nakamura, T., & Akazawa, C. (2017). Differentiation of Oligodendrocyte Precursor Cells from Sox10-Venus Mice to Oligodendrocytes and Astrocytes. Sci Rep, 7(1), 14133. cccccc10.1038/s41598-017-14207-0

Szewczyk, L. M., Lipiec, M. A., Liszewska, E., Meyza, K., Urban-Ciecko, J., Kondrakiewicz, L., Goncerzewicz, A., Rafalko, K., Krawczyk, T. G., Bogaj, K., Vainchtein, I. D., Nakao-Inoue, H., Puscian, A., Knapska, E., Sanders, S. J., Jan Nowakowski, T., Molofsky, A. V., & Wisniewska, M. B. (2024). Astrocytic beta-catenin signaling via TCF7L2 regulates synapse development and social behavior. Mol Psychiatry, 29(1), 57–73. 10.1038/s41380-023-02281-y

Talley, M. J., Nardini, D., Ehrman, L. A., Lu, Q. R., & Waclaw, R. R. (2023). Distinct requirements for Tcf3 and Tcf12 during oligodendrocyte development in the mouse telencephalon. Neural Dev, 18(1), 5. 10.1186/s13064-023-00173-z

Tsyganov, K., Perry, A., Archer, S., & Powell, D. (2018). MonashBioinformaticsPlatform/RNAsik-pipe: JOSS ready In (Version 1.5.5)

Wang, W., Feng, Y., Aimaiti, Y., Jin, X., Mao, X., & Li, D. (2018). TGFbeta signaling controls intrahepatic bile duct development may through regulating the Jagged1-Notch-Sox9 signaling axis. J Cell Physiol, 233(8), 5780–5791. 10.1002/jcp.26304

Wang, Y., Li, G., Stanco, A., Long, J. E., Crawford, D., Potter, G. B., Pleasure, S. J., Behrens, T., & Rubenstein, J. L. (2011). CXCR4 and CXCR7 have distinct functions in regulating interneuron migration. Neuron, 69(1), 61–76. 10.1016/j.neuron.2010.12.005

Wolf, F. A., Angerer, P., & Theis, F. J. (2018). SCANPY: large-scale single-cell gene expression data analysis. Genome Biol, 19(1), 15. 10.1186/s13059-017-1382-0

Wu, T., Hu, E., Xu, S., Chen, M., Guo, P., Dai, Z., Feng, T., Zhou, L., Tang, W., Zhan, L., Fu, X., Liu, S., Bo, X., & Yu, G. (2021). clusterProfiler 4.0: A universal enrichment tool for interpreting omics data. Innovation (Camb), 2(3), 100141. 10.1016/j.xinn.2021.100141

Yu, G., & He, Q. Y. (2016). ReactomePA: an R/Bioconductor package for reactome pathway analysis and visualization. Mol Biosyst, 12(2), 477–479. 10.1039/c5mb00663e

Yun, K., Fischman, S., Johnson, J., Hrabe de Angelis, M., Weinmaster, G., & Rubenstein, J. L. (2002). Modulation of the notch signaling by Mash1 and Dlx1/2 regulates sequential specification and differentiation of progenitor cell types in the subcortical telencephalon. Development, 129(21), 5029–5040. 10.1242/dev.129.21.5029

Zhang, M. Z. (2019). Notch signaling is essential in collecting duct epithelial cell fate determination during development and maintenance of cell type homeostasis in adult. Ann Transl Med, 7(Suppl 8), S376. 10.21037/atm.2019.12.121

Zhou, Q. P., Le, T. N., Qiu, X., Spencer, V., de Melo, J., Du, G., Plews, M., Fonseca, M., Sun, J. M., Davie, J. R., & Eisenstat, D. D. (2004). Identification of a direct Dlx homeodomain target in the developing mouse forebrain and retina by optimization of chromatin immunoprecipitation. Nucleic Acids Res, 32(3), 884–892. 10.1093/nar/gkh233

Zhou, Y., Zhou, B., Pache, L., Chang, M., Khodabakhshi, A. H., Tanaseichuk, O., Benner, C., & Chanda, S. K. (2019). Metascape provides a biologist-oriented resource for the analysis of systems-level datasets. Nat Commun, 10(1), 1523. 10.1038/s41467-019-09234-6

