## Supplementary Figures for "The DLX/Notch axis is necessary for spatiotemporal regulation of neural cell fate"

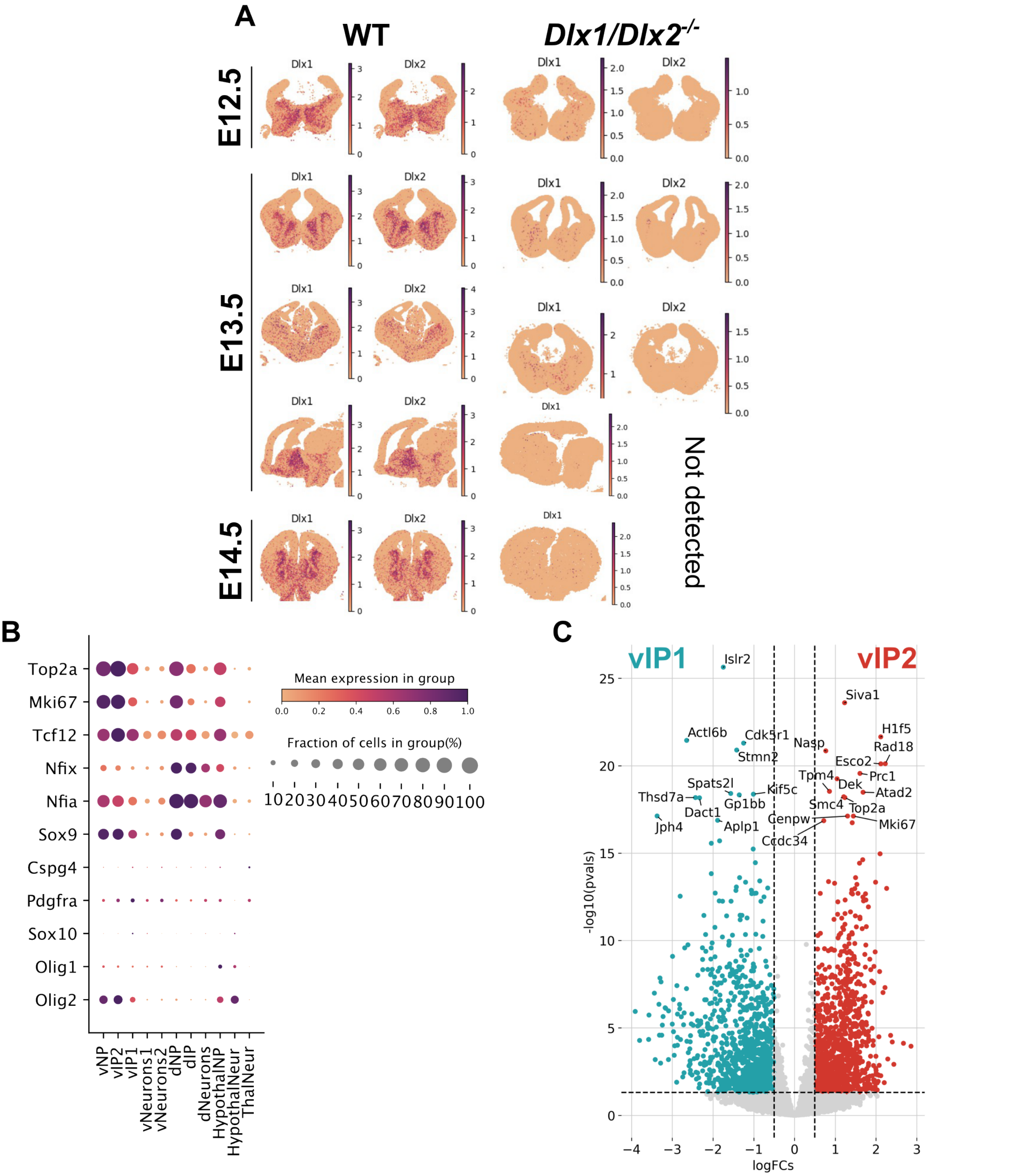


Leung et al., 2025 Figure S1

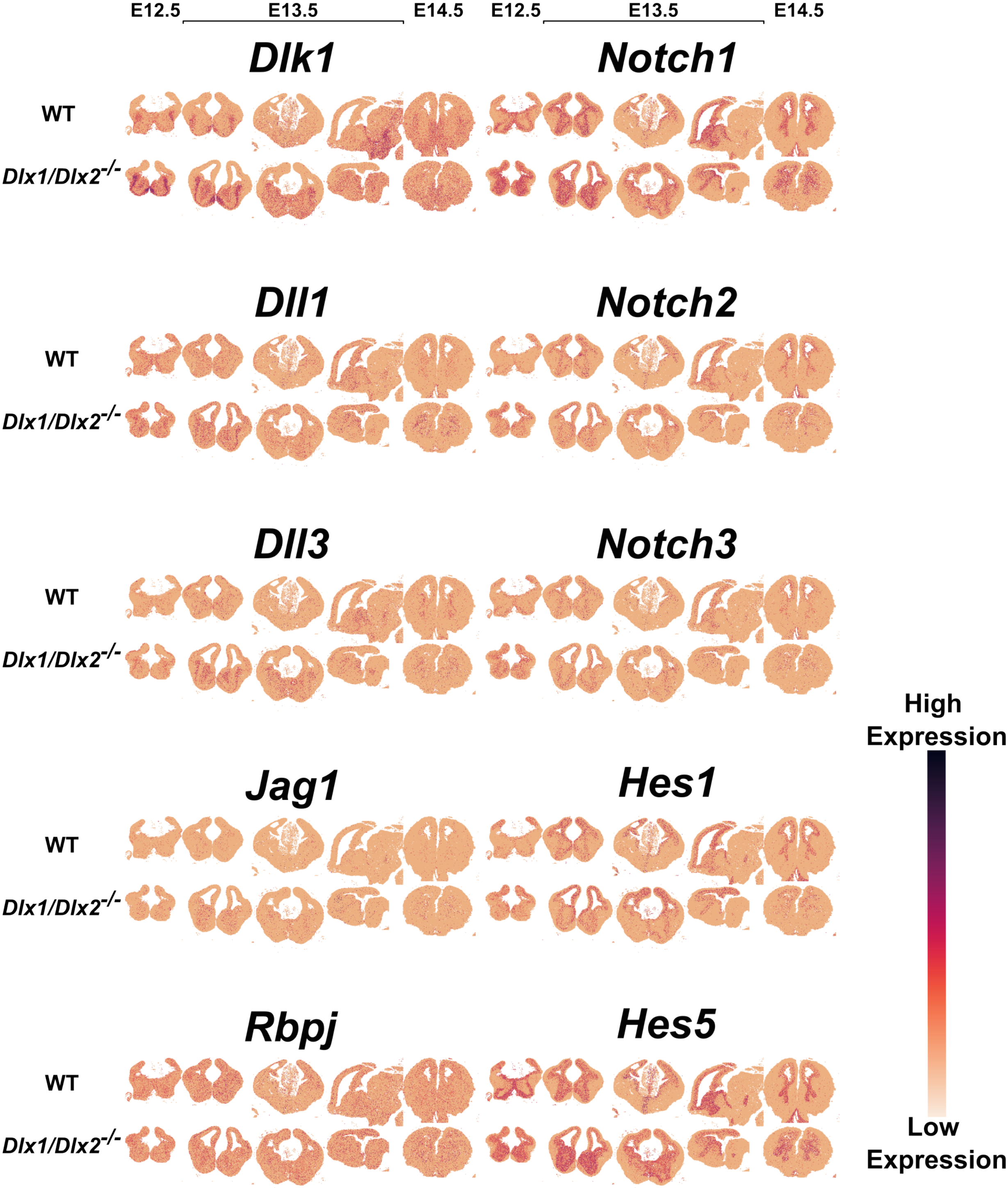


Leung et al., 2025 Figure S2

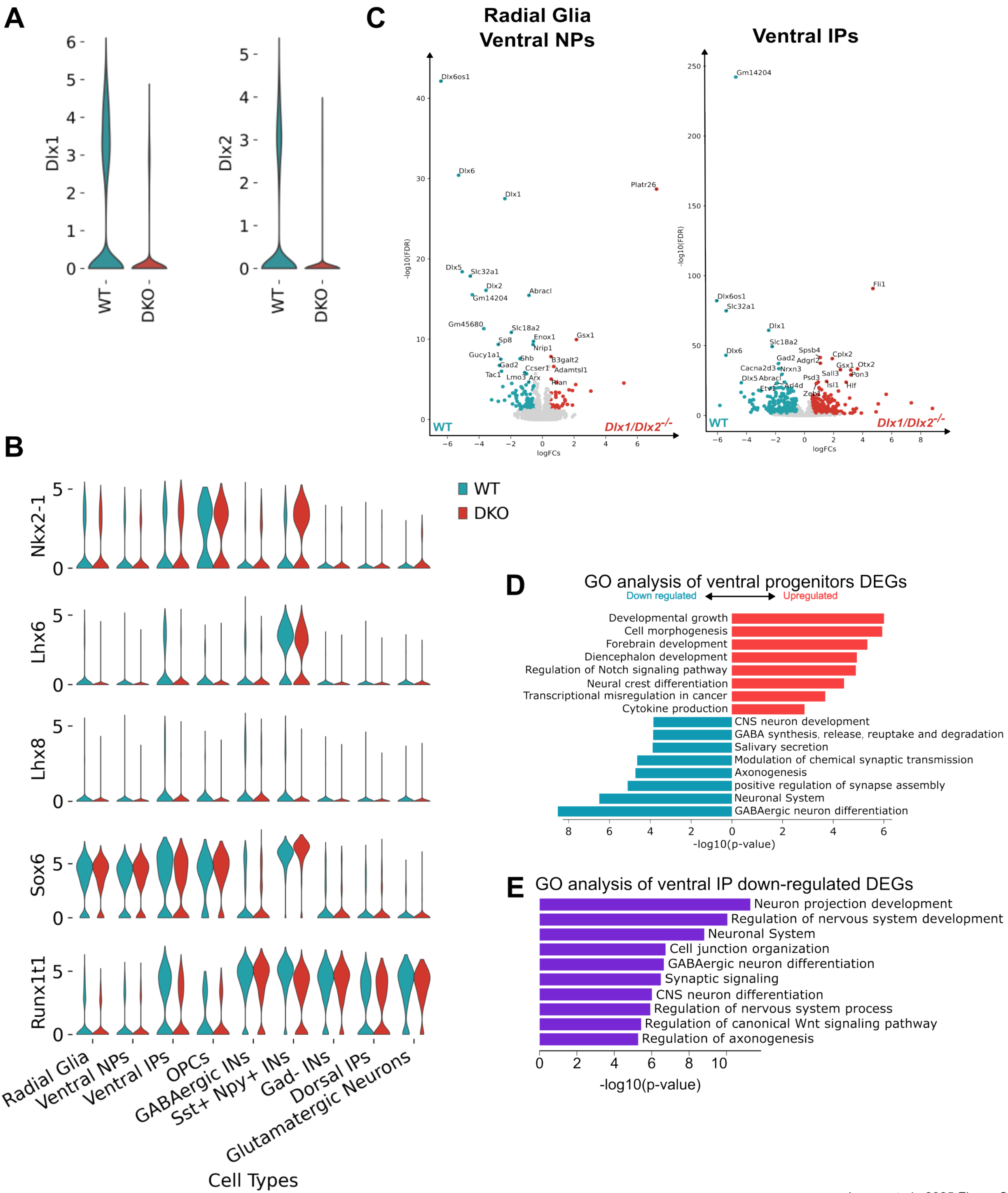


Leung et al., 2025 Figure S3

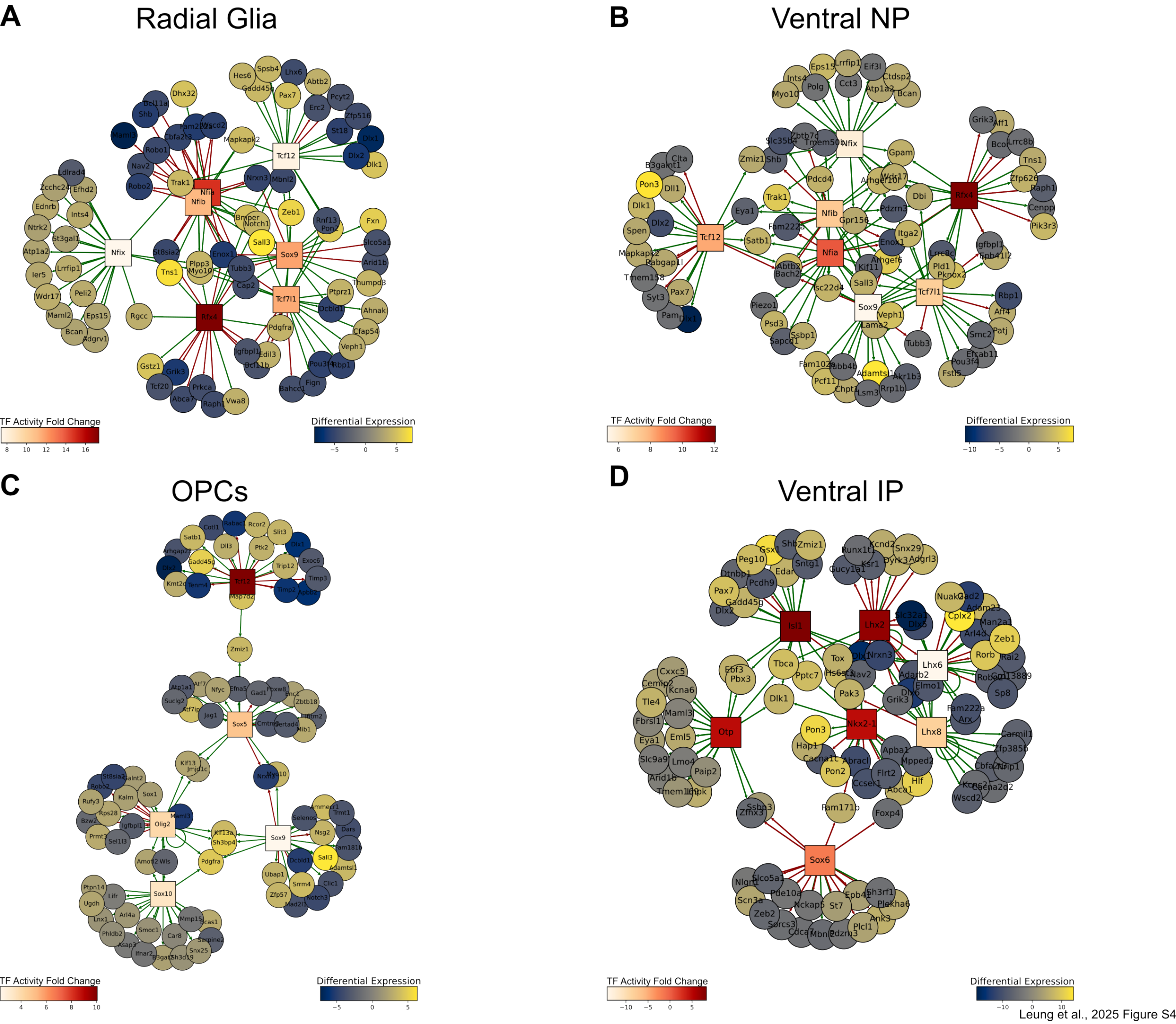


Leung et al., 2025 Figure S4

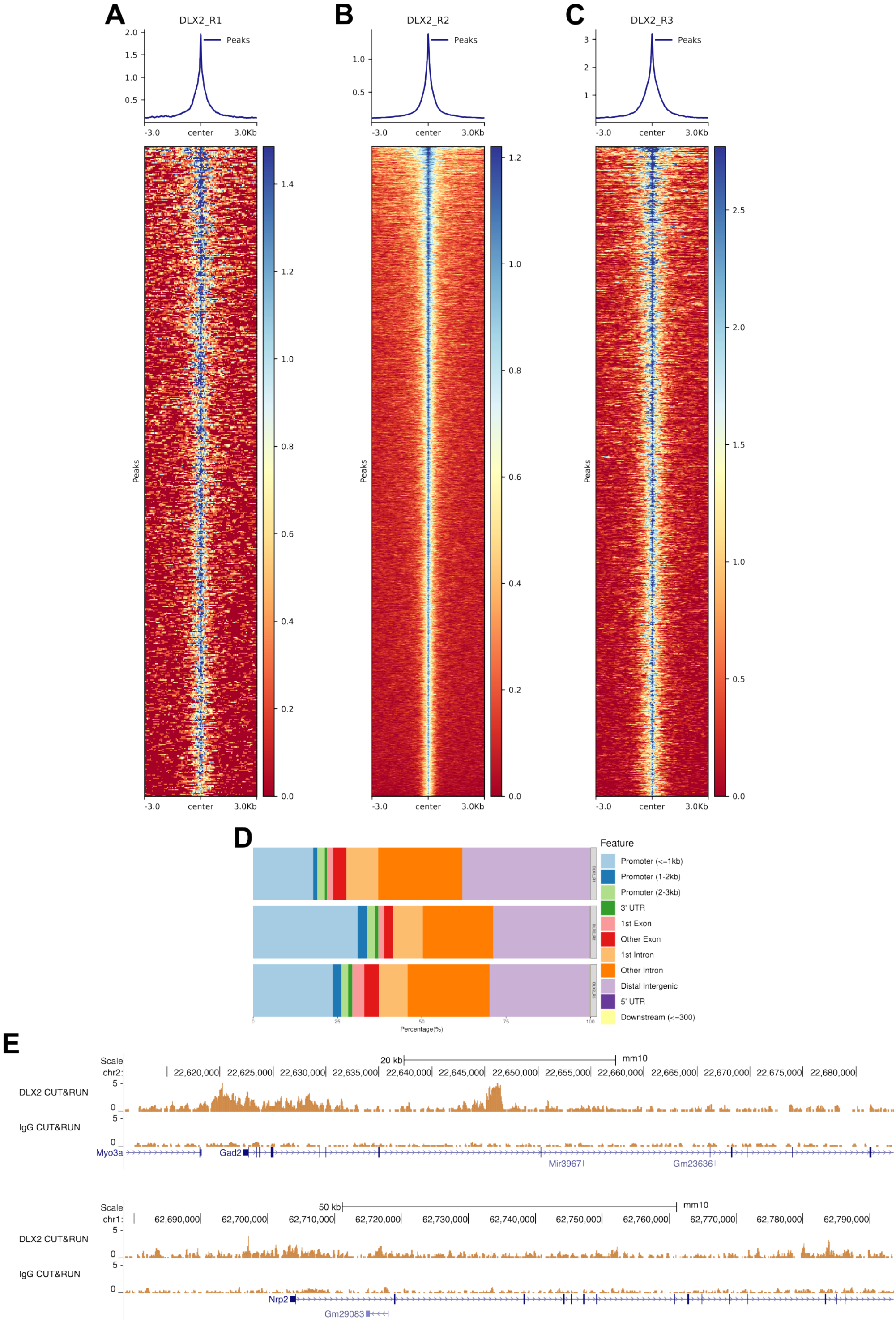

A
DLX2_R1
Peaks
-3.0
center
3.0Kb
B
DLX2_R2
C
DLX2_R3
D
Feature
Promoter (<=1kb)
Promoter (1-2kb)
Promoter (2-3kb)
3' UTR
1st Exon
Other Exon
1st Intron
Other Intron
Distal Intergenic
5' UTR
Downstream (<=300)
Percentage(%)
E
Scale
chr2:
20 kb
mm10
DLX2 CUT&RUN
IgG CUT&RUN
Myo3a
Gad2
Mir3967
Gm23636
chr1:
50 kb
Nrp2
Gm29083

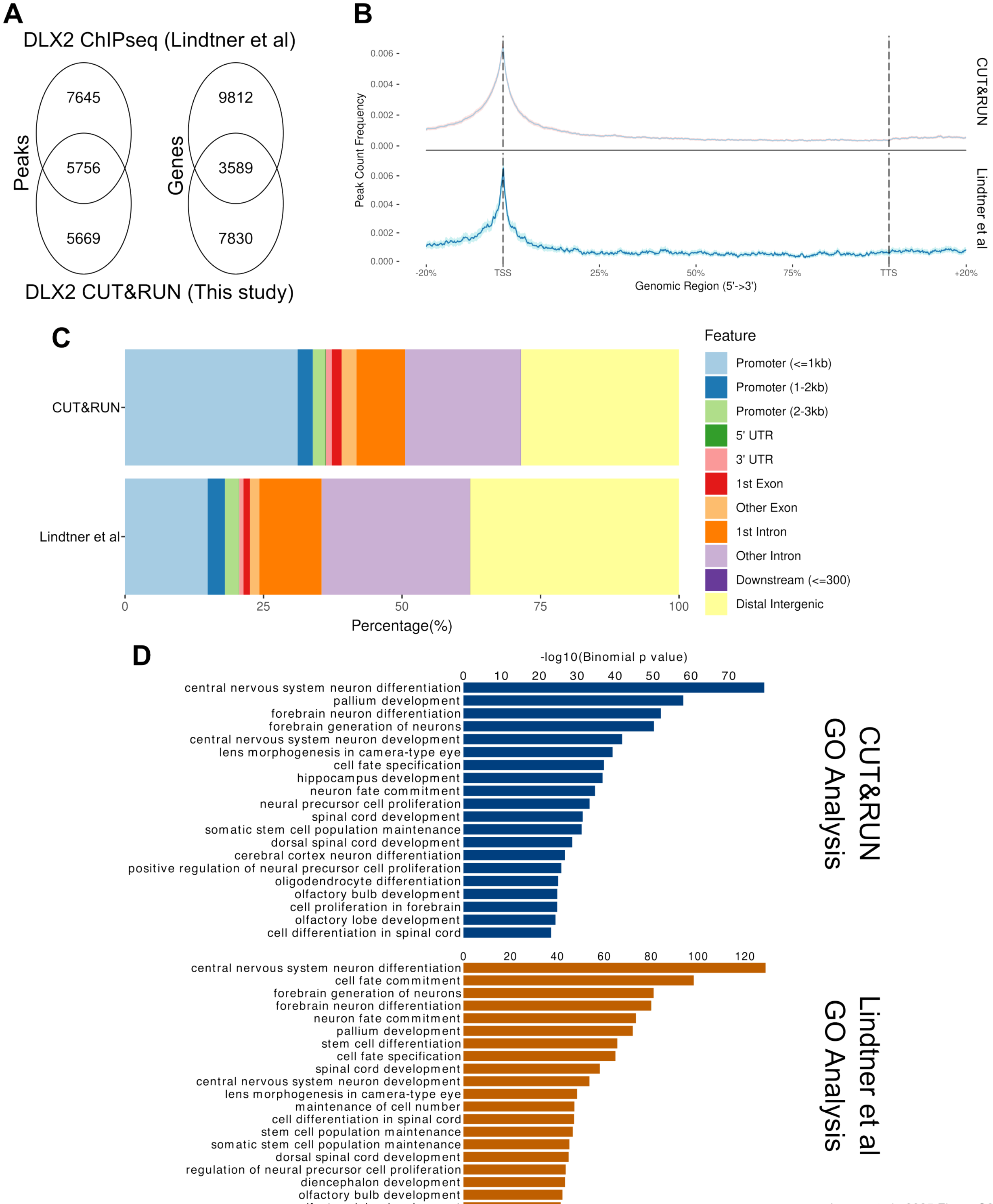
A
DLX2 ChIPseq (Lindtner et al)
Peaks
7645
5756
5669
Genes
9812
3589
7830
DLX2 CUT&RUN (This study)
B
Peak Count Frequency
0.006
0.004
0.002
0.000
CUT&RUN
Lindtner et al
-20%
TSS
25%
50%
75%
TTS
+20%
Genomic Region (5'->3')
C
CUT&RUN
Lindtner et al
0
25
50
75
100
Percentage(%)
Feature
Promoter (<=1kb)
Promoter (1-2kb)
Promoter (2-3kb)
5' UTR
3' UTR
1st Exon
Other Exon
1st Intron
Other Intron
Downstream (<=300)
Distal Intergenic
D
-log10(Binomial p value)
0 10 20 30 40 50 60 70
central nervous system neuron differentiation
pallium development
forebrain neuron differentiation
forebrain generation of neurons
central nervous system neuron development
lens morphogenesis in camera-type eye
cell fate specification
hippocampus development
neuron fate commitment
neural precursor cell proliferation
spinal cord development
somatic stem cell population maintenance
dorsal spinal cord development
cerebral cortex neuron differentiation
positive regulation of neural precursor cell proliferation
oligodendrocyte differentiation
olfactory bulb development
cell proliferation in forebrain
olfactory lobe development
cell differentiation in spinal cord
CUT&RUN GO Analysis
0 20 40 60 80 100 120
central nervous system neuron differentiation
cell fate commitment
forebrain generation of neurons
forebrain neuron differentiation
neuron fate commitment
pallium development
stem cell differentiation
cell fate specification
spinal cord development
central nervous system neuron development
lens morphogenesis in camera-type eye
maintenance of cell number
cell differentiation in spinal cord
stem cell population maintenance
somatic stem cell population maintenance
dorsal spinal cord development
regulation of neural precursor cell proliferation
diencephalon development
olfactory bulb development
olfactory lobe development
Lindtner et al GO Analysis

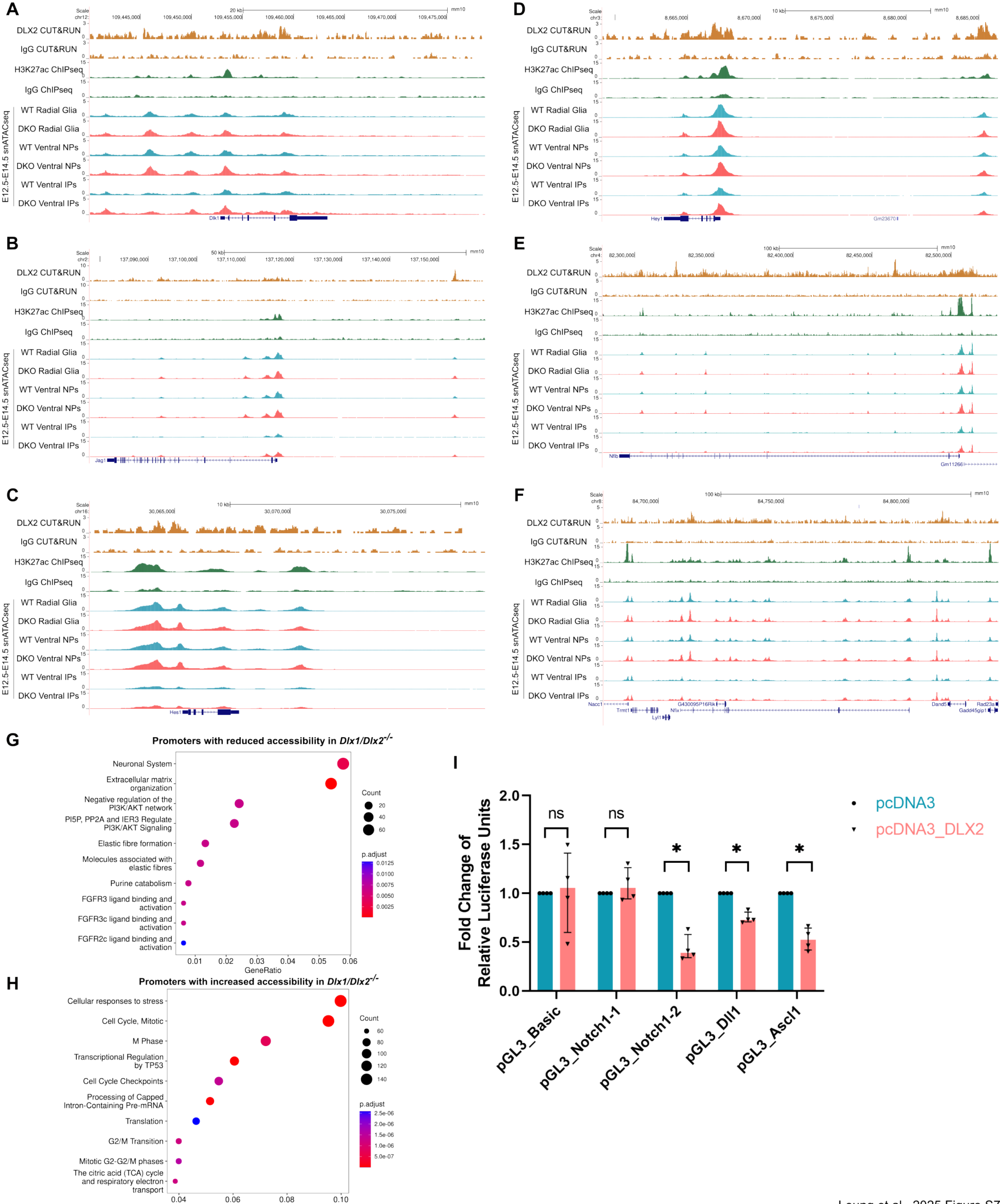


Leung et al., 2025 Figure S7
